# Standard benchmarking inflates gene-prioritization accuracy estimates

**DOI:** 10.64898/2026.11.24.768897

**Authors:** Matevž Kovačič

## Abstract

Benchmark results tell clinics which gene-prioritization system to trust and the field what counts as progress. Standard benchmarking practice — testing each system using its latest knowledge databases — can inflate system accuracy estimates and invite data leakage. We formalize the Fairbench protocol: testing may use only knowledge available strictly before the patient’s assessment. Against prepublication snapshots dated by case report, on one public case collection, standard benchmarking inflated Top-1 accuracy by 40.5–80.5 percentage points across Exomiser, LIRICAL, Phrank and Phen2Disease. In every system, over half the tested patients had a causal-gene phenotype association whose only recorded source was their case publication. Reassessing three systems’ published benchmarks with reproduced or reconstructed references showed the same direction in all five comparisons. Retesting undiagnosed patients with the latest knowledge database is good care. But benchmark results are read as performance at the patient’s assessment, and Fairbench is a necessary data-hygiene condition for that reading.

---

Gene-prioritization systems rank the genes a clinician should investigate in a patient with a suspected rare disease [1], and are in clinical use: Exomiser supplied the phenotype-based prioritization in the 100,000 Genomes Project’s rare-disease pilot [2]. Their benchmark results carry real decisions: they inform which system a clinic deploys, and the field reads them as progress [3]. Behind that choice are patients whose search for the causal gene these systems can shorten or prolong. Those decisions are only as good as the measurement — a benchmark claiming historical performance must estimate how well a system could have served the patient at the time of the assessment, not how well it retrieves answers the field has since written down.

When those later answers derive from the patient’s own case report, this is data leakage: the case under test supplied part of the knowledge used to solve it. Later knowledge has a legitimate use on today’s patient — reanalysing an undiagnosed case is good care; what misleads is presenting the later run’s accuracy as performance at the earlier assessment.

Earlier evaluations have shown why the timing and source of diagnostic knowledge matter for interpreting benchmark accuracy. Leakage of evaluation-linked information into the knowledge a model uses is a recognized failure mode across machine-learning evaluation [4, 5], and community assessments in biology control it with explicit time cutoffs [6]. Börnigen and colleagues, testing gene-prioritization systems on associations not yet in their databases, warned that retrospective evaluations can be overly optimistic [7]; a later study found one system weaker on newly discovered associations than in cross-validation [8]; historical databases have also evaluated simulated patients whose associations were discovered after release [9]. Overlap between training data and evaluation sets produced over-optimistic comparisons of variant-effect predictors [10], and guiding questions for avoiding leakage in biological machine learning are now in print [11].

Most directly, two preprint studies deleted case-linked knowledge from fixed releases of the systems’ knowledge databases and showed that scores depend on it [12, 13].

Here we measure the complementary quantity: how benchmark accuracy changes across dated releases of each system’s knowledge databases when the same patients are evaluated by the same ranking implementations. We do this across four published diagnostic systems, from variant-aware to phenotype-only prioritization. We examine dated knowledge-database effects, candidate availability and documented source reuse, each in its own analysis; test the documented reuse by removing the sole-source associations in one system at a fixed release; and reassess published benchmark configurations.

The common retrospective design runs every patient case only against each system’s latest data release [14–16] — a single snapshot of the diagnostic system’s knowledge database. That snapshot postdates all but the most recent case reports. To prevent temporal contamination, we formalize the Fairbench protocol: for each patient, the system must prioritize candidate genes using only knowledge publicly available strictly before the prioritization being evaluated. Fig. 1 contrasts the protocol with standard practice.

Our experiments show that using later knowledge databases substantially increases the measured accuracy of the systems being benchmarked. We evaluated Exomiser, LIRICAL, Phrank and Phen2Disease [15–18]. For each patient, we compared results obtained with each system’s most recent database versions released strictly before the patient’s case report with results obtained using later versions. Patient inputs, ranking implementations and settings remained fixed. Absence from a resource does not mean the association was undiscovered, because curation lags publication.

**Fig. 1.**
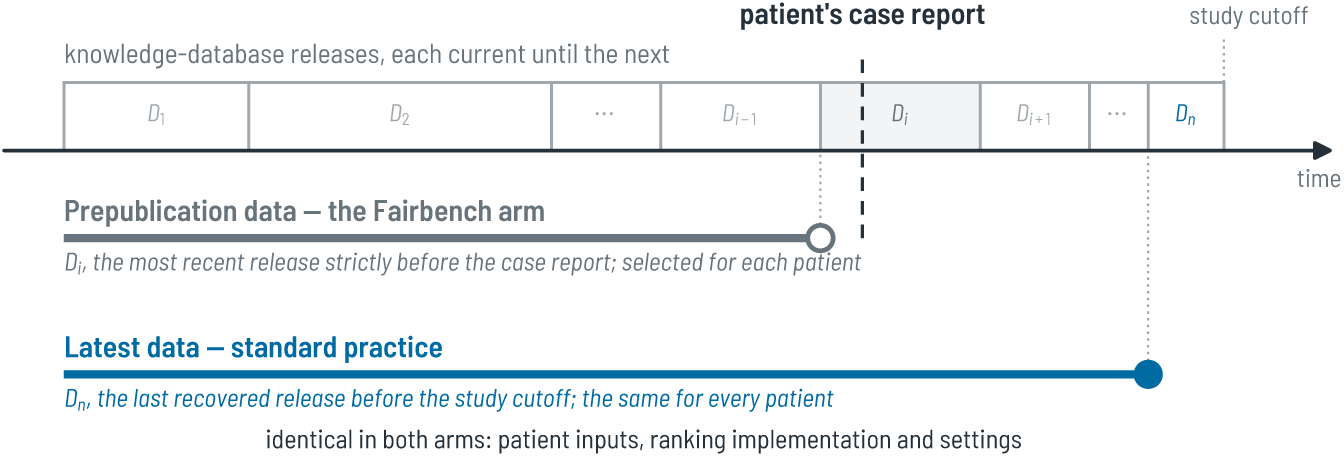
The Fairbench protocol against standard benchmarking practice. Knowledge databases issue releases at irregular intervals (*D*_1_*, . . . , D_n_*), each release current until the next. The figure shows this study’s implementation of the protocol, with case publication as the practical reference for the prioritization being evaluated. Standard benchmarking runs each system with the latest data: the last recovered release before the study cutoff (*Dn*, filled circle), which postdates the illustrated patient’s case report. The Fairbench arm instead uses the prepublication data: each knowledge database’s most recent release issued strictly before that patient’s case report (*D_i_*, open circle).

The later knowledge databases contained findings sourced from the benchmarked cases’ own publications, in all four systems, with native replay in Exomiser and LIR-ICAL confirming their participation in scoring. Removing the associations supported only by each patient’s publication sharply reduced Phrank’s accuracy at the fixed final release.

We also reassessed published benchmark evaluations of Exomiser, LIRICAL and Phen2Disease: replacing the knowledge databases with versions released before each patient’s case report lowered accuracy in all five comparisons; four of the five Top-1 95% confidence intervals excluded zero.

## 1 Results

We first asked how much ranking accuracy changes when the same patients and algorithms are evaluated with knowledge-database releases from different dates. A patient entered a system’s comparison only when a release predating their case report was recoverable; the resulting cohorts, drawn from a common catalogue of 10,377 published cases in Phenopacket Store [19], are given with their earliest usable releases in Table 3 (Online Methods). The 1,090 patients eligible for all four systems form the shared cohort.

### 1.1 Later knowledge-database releases raise causal-gene ranking accuracy

Changing the supplied knowledge databases increased how often the reported causal gene ranked first (Top-1 accuracy) by 40.5–80.5 percentage points across the four systems (Fig. 2a). Patient inputs, algorithm implementations and configurations remained unchanged (Supplementary Table S1).

Large increases in gene-prioritization accuracy remained when all four systems evaluated the same 1,090 patients (Fig. 2a–c, shared-cohort rows), so the finding held with patient membership matched across systems. In both cohort comparisons, later knowledge databases also improved Top-3 and Top-10 accuracy: the causal gene appeared among the first three or ten candidates more often.

**Fig. 2.**
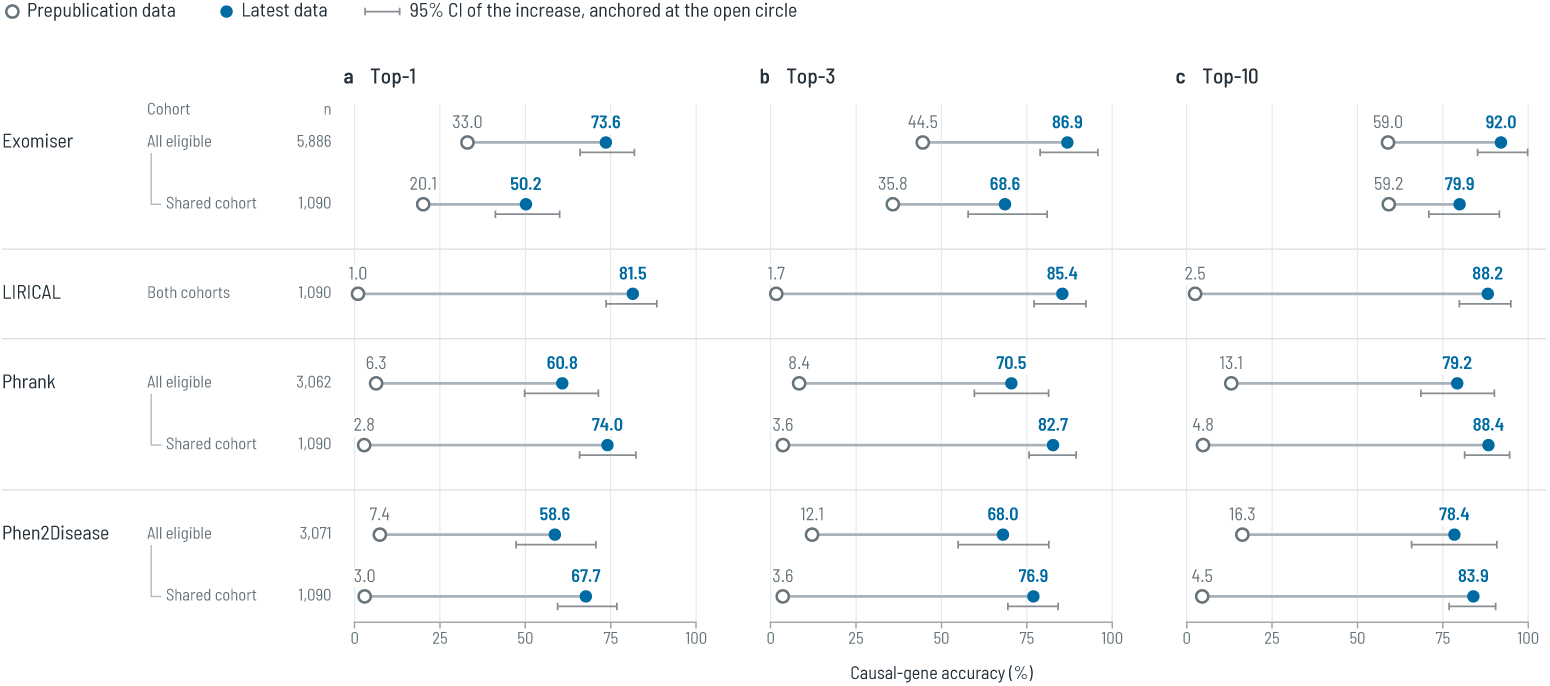
Causal-gene ranking accuracy with prepublication versus latest knowledge-database releases. **a–c**, Top-1, Top-3 and Top-10 accuracy. Within each system, the upper row gives all its eligible cases and the lower row the shared cohort: the same 1,090 patients in every system (cohort sizes: Table 3). LIRICAL’s eligible cohort is the shared cohort, so it appears once. Open circles give accuracy with the prepublication releases, each patient’s most recent release before their case report; filled circles with each system’s latest recovered release (Online Methods). The bracket below each pair is the 95% publication-cluster confidence interval of the increase, anchored at the open circle. Increases are calculated before rounding.

Successive releases show increases, plateaus and occasional decreases within fixed patient groups (Extended Data Fig. 1). These illustrative groups are concentrated in particular publications; they are not representative samples of clinical referrals. Supplementary Figs. S1 and S2 show all available pairs and the fixed-group Top-3 and Top-10 trajectories.

The size of the increase depends on which release ends the comparison: holding January’s already-evaluated patients fixed, most of each phenotype-only system’s increase was in place by 16 February 2026 (Supplementary Table S2). A final score alone obscures this concentration in particular releases.

### 1.2 Knowledge-database updates change both candidate availability and ranking

A higher score can reflect the return of a previously absent causal gene or a better rank for one already returned. In Exomiser, almost the entire increase occurred among genes returned at both endpoints; in the other three systems, newly returned genes contributed most of it (Supplementary Table S3). These contributions use whole-cohort denominators. Candidate presence does not establish a known disease association or an unchanged set of competing genes.

The candidate shifts have a resource-level counterpart. For a third of Exomiser’s patients, two thirds of LIRICAL’s and about half of Phrank’s and Phen2Disease’s, the baseline release held no phenotype association for the causal gene (for Exomiser, counted in its human phenotype data only). Ranking such a gene is the discovery setting. There, the three phenotype-only systems have zero prepublication Top-1 accuracy; Exomiser has 5.6%, through its other evidence channels. At each system’s latest knowledge-database release the same patients score 58.9–84.3%, and their gains contribute 18.9–56.0 points of each system’s overall increase. Little of this lies beyond a removal control’s reach: in every system, only a small fraction of these patients ends with no association supported by their own paper alone, and theirs are the only gains such a control cannot reach (patient counts, per-system values and intervals: Supplementary Table S5).

The large increases did not require a distant prepublication baseline: limiting the baseline-to-publication interval to 90 days retained changes of 34.5–80.0 points, and equal weighting of genes, publications or recorded disease-label sets, and exclusion of recorded missing or mapped phenotype terms, also retained large effects (Supplementary Tables S7, S8 and S9). Further sensitivity analyses cluster the variant-aware results by background donor and split established associations by the basis of their dating evidence (Supplementary Tables S10 and S12). These checks address specific alternatives; they do not establish clinical representativeness or complete historical compatibility.

### 1.3 Knowledge accumulation includes findings from benchmark publications

Evidence about a tested patient can reach a knowledge database by several routes, including laboratory data and related patients; we examined one traceable route: whether the knowledge databases incorporated findings from the publications describing the evaluated patients. The audit does not assess every possible information source or establish independence between related patients.

The knowledge databases incorporated findings from papers describing the very patients being benchmarked. These findings supplied phenotype associations used to rank those patients’ causal genes. In Exomiser, these phenotype associations entered the comparison between a patient’s phenotypes and human disease profiles; because the exported profiles retain no citations, this linkage is inferred by matching the consumed associations to the database provider’s declared upstream annotations (Online Methods). In LIRICAL, these phenotype associations entered calculations of how strongly the patient’s findings supported a candidate disease. In Phrank and Phen2Disease, these phenotype associations formed part of the gene–phenotype profiles used to rank candidate genes.

For more than half the patients tested in each benchmark, the later databases contained phenotype associations whose only recorded support came from the paper describing those patients. Corpus and knowledge databases draw on the same clinical literature: the HPO annotations cite case publications as sources [20], and Phenopacket Store assembles its cases from such publications. Table 1 compares pre-publication and later-database accuracy for this patient group. It also reports how much the group’s score changes contributed to the overall benchmark increase.

Native scoring checks established that source-linked associations reach the scoring calculations in Exomiser and LIRICAL (supplementary section “Publication sources and native scoring”). Patients with these documented links also accounted for much of the observed increase (Table 1); other database updates could have improved the same patients’ rankings, so the table does not establish causation.

**Table 1.** Top-1 results for patients with at least one causal-gene phenotype association supported exclusively by their own paper in the audited records. Patients: how many of each system’s cohort are in this group. Historical and Final: the group’s Top-1 accuracy (%) with the prepublication and the latest releases. The whole-cohort contribution (percentage points, with the 95% publication-cluster confidence interval) uses the full cohort denominator and describes observed changes, not causal effects.

| System | Patients | Historical | Final | Whole-cohort contribution (95% CI) |
|---|---|---|---|---|
| Exomiser | 3,347 | 21.75 | 76.07 | 30.89 (23.96–38.83) |
| LIRICAL | 1,006 | 0.40 | 86.88 | 79.82 (71.52–86.83) |
| Phrank | 2,466 | 4.98 | 72.81 | 54.63 (44.23–64.94) |
| Phen2Disease | 2,475 | 3.71 | 67.63 | 51.51 (40.87–63.11) |

We then tested this dependence directly in one system. Phrank consumes a resource that records support for each row, so the audited sole-source classification maps onto an executable deletion, verified file for file against the audit. At the fixed final release, we removed the phenotype associations supported only by each patient’s own publication — the propagated associations of every gene and every disease that publication alone supports, not only the patient’s gene or diagnosis — and rebuilt Phrank’s ranking model for each of the 1,690 source publications (Online Methods). The removal reduced Phrank Top-1 accuracy by 50.83 percentage points (95% CI 40.80 to 60.44), from 60.81% to 9.97%, in the 3,062 patients of the temporal experiment’s Phrank cohort, and by 29.09 points (24.89 to 33.43), from 52.51% to 23.42%, across all 9,588 catalogued patients with an NCBI Gene identifier for the causal gene (Extended Data Table 1). Patients whose causal gene loses all its associations remain in the denominators with zero credit. This deletion effect is measured at one release; it is not a share of the dated increase.

### 1.4 The system authors’ published benchmarks change with the knowledge-database release

We retested the system authors’ original patients to ask whether the knowledge-database release used in testing each patient changes their published benchmark results, reusing their patient inputs and software and recovering the accompanying resources as fully as public materials allowed. Phen2Gene’s algorithm was excluded: rebuilding its knowledge resources did not reproduce its published term tables, so differences could not be attributed to the knowledge-database release alone.

We compared reproduced or reconstructed reference runs with runs on the same patients using resource snapshots released strictly before each patient’s case report (Table 2). The comparisons retained 2,744 of 4,677 Exomiser cases, 313 of 384 LIR-ICAL cases and 615 of 2,051 Phen2Disease source entries, with the algorithm and patient inputs fixed within each comparison. The reproduced Exomiser reference is the PhEval phenotype-only arm, not the variant-aware configuration in our main experiment (the same release and mode as a recent comparison with large language models [3], which we do not rerun).

**Table 2.** Accuracy on cases from published benchmarks, using knowledge databases released before each patient’s case report versus the reproduced or reconstructed benchmark reference. Each comparison uses the same cases and scoring rules. Accuracy is expressed as a percentage; bold increases and parenthesized 95% confidence intervals are percentage points. LIRICAL ranks diseases; the other systems rank genes. **^ψ^**Exomiser is the reproduced PhEval phenotype-only arm; LIRICAL uses recovered 2022 resources, and Phen2Disease uses reconstructed resources. Phen2Disease’s row labels name the source patient collections, assembled for LIRICAL, Phen2Gene and PhenoApt [21]; each is analysed separately.

| System and comparison | Cases | Before<br>case report | Benchmark<br>reference | Increase<br>(95% CI) |
|---|---|---|---|---|
| *Top-1* |  |  |  |  |
| Exomiser<sup>†</sup> | 2,744 | 3.6 | 28.4 | **24.9** (15.9–38.9) |
| LIRICAL (2022 reference) | 313 | 10.5 | 22.0 | **11.5** (5.7–17.4) |
| Phen2Disease: LIRICAL | 292 | 12.7 | 37.2 | **24.6** (17.2–31.8) |
| Phen2Disease: Phen2Gene | 144 | 3.5 | 30.9 | **27.4** (10.4–43.3) |
| Phen2Disease: PhenoApt | 179 | 21.0 | 23.1 | **2.1** (-2.4–6.7) |
| *Top-3* |  |  |  |  |
| Exomiser<sup>†</sup> | 2,744 | 6.7 | 36.2 | **29.5** (18.2–45.1) |
| LIRICAL (2022 reference) | 313 | 19.5 | 38.0 | **18.5** (12.5–24.5) |
| Phen2Disease: LIRICAL | 292 | 20.1 | 46.7 | **26.6** (18.7–34.3) |
| Phen2Disease: Phen2Gene | 144 | 5.8 | 39.6 | **33.8** (16.9–49.2) |
| Phen2Disease: PhenoApt | 179 | 28.9 | 36.4 | **7.5** (2.4–13.0) |
| *Top-10* |  |  |  |  |
| Exomiser<sup>†</sup> | 2,744 | 14.5 | 43.3 | **28.8** (18.7–44.0) |
| LIRICAL (2022 reference) | 313 | 27.2 | 49.8 | **22.7** (15.6–29.5) |
| Phen2Disease: LIRICAL | 292 | 30.7 | 61.4 | **30.8** (21.7–39.2) |
| Phen2Disease: Phen2Gene | 144 | 7.7 | 48.6 | **40.9** (23.4–56.0) |
| Phen2Disease: PhenoApt | 179 | 41.1 | 48.7 | **7.7** (1.9–13.7) |

Four of the five Top-1 differences exclude zero; the exception is Phen2Disease on PhenoApt cases (2.1 points, *−*2.4 to 6.7).

Exomiser reproduced the deposited scores at four-decimal precision and worsttie ranks for 4,676 of 4,677 cases (the one mismatch is excluded and documented in the supplement). LIRICAL’s original 2019 annotations and Phen2Disease’s similarity matrix were unavailable: those comparisons measure changes from reconstructed references, not corrections to the published headline accuracies. Not every original case could be retested — mainly for missing publication dates or unavailable snapshots — so the changes describe only cases tested under both conditions (exclusions and reconstruction checks: Supplementary Notes, “Published-benchmark reconstruction”; Top-3 and Top-10: Supplementary Fig. S3).

## 2 Discussion

Running the same gene-prioritization system benchmark with a different release of the system’s knowledge databases can change the benchmark’s conclusions, with the system’s ranking algorithm untouched. Across the four systems tested in this paper, the use of later knowledge-database releases substantially raised system accuracy on the same patient inputs. The increases stayed large on the shared cohort, our patients every system can process.

Our experiments estimate the change in a system’s gene-prioritization accuracy between two settings. In one, the system tests each patient using only the system’s last knowledge database predating the patient’s publication; in the other — the standard benchmarking practice — it tests the patient with its latest, most complete knowledge database. Everything else, mainly the ranking algorithm, is held constant. The measured increase in the system’s accuracy is inflation: by that amount, standard benchmarking overstates the accuracy the system could have delivered when the case was reported.

Within each system’s paired comparison between prepublication and latest knowledge-database use, the observed accuracy difference is attributable to that change and nothing else. A latest-snapshot knowledge-database benchmark uses the most complete one: evidence from the tested cases’ own publications can be in it. This inflation is a bias, not sampling noise: testing more patients does not reduce it, because every added patient is scored against the same too-complete knowledge database. More data never cures contamination; related cases sitting in both the knowledge database and the test set are another example. Testing each patient with their prepublication knowledge databases makes the case out of sample with respect to later releases; it does not make the selected states independent of one another, or certify all patient observations and fixed components as historically available. Individual knowledge-database releases can still influence many cases, as the release trajectories and endpoint comparisons show. The 95% intervals cluster patients by source publication — one publication can describe many patients — and quantify sampling variation only, not the choice of knowledge-database states.

Using accumulated knowledge in the system’s latest knowledge database has its place and is invaluable for patient care: discoveries from a previous patient can help solve a new case or reanalyse an unresolved one [22]. In contrast, benchmarking that claims historical performance is a measurement: it asks how well a system could have served an unseen patient at the time of assessment.

Standard benchmarking retests patients whose discoveries have since entered the knowledge database, so it scores the system partly on cases the database already contains — ranked with associations linked to the benchmark publication — and partly on cases where knowledge from the patient’s future may still intrude.

Our experiments measure the paired contrast; our four-system audit traces one of its leak pathways: whether the knowledge used to score a patient came from that patient’s own publication. The consumed phenotype resources carry source records — directly or, for Exomiser, through its declared upstream annotation release — and we read them for every evaluated patient. We measured the extent of the documented overlap: how many patients are affected, and how much benchmark credit occurs among cases with publication-source overlap. For each association counted in that overlap, the audited record documents the case publication as the sole source and no earlier one. How much any single association changed the increase we cannot say; the overlap itself is documented in the consumed resources’ own records.

In related work, two preprint studies removed case-linked knowledge from a fixed release of the system’s knowledge database. One dropped each case’s gold-profile terms supported only by its own publication (leave-source-out) [12]. The other deleted the disease–phenotype annotations a source publication alone supports (leave-onepublication-out) [13]. Both showed that scores depend on these entries. Neither reported how later releases of the system’s knowledge database change measured accuracy. We did both kinds of experiment. The dated comparisons measure the change across releases: this paper’s four systems, per-patient resource dates, three reassessed published benchmarks. The removal repeats the deletion test in Phrank, under a sole-source rule [12, 13]: it deletes the phenotype associations the tested patient’s own publication alone supports, across every disease and gene, and rebuilds Phrank’s model for each publication. It ran on the temporal experiment’s patients, in its implementation, at its final knowledge-database release. For every patient, the two contrasts therefore stand side by side. They mostly agree in direction (Supplementary Table S15). Agreement is all we claim; neither quantity is a share of the other. Their LIRICAL deletion is phenotype-only; it corroborates the phenotype channel, not the variant-aware LIRICAL configuration tested here. The removal design is theirs. The dated, per-patient comparison, and its pairing with a removal on the same patients, are this study’s addition.

Retesting the system authors’ own benchmark patients shows the same: accuracy changes with the system’s knowledge-database release used in testing each patient. The PhEval phenotype-only Exomiser reference was reproduced, allowing a paired comparison on its historically retestable cases. LIRICAL and Phen2Disease required reconstructed references because some original resources were unavailable. Their results establish effects in the recovered setups; they cannot be subtracted from published headline accuracies, or used to infer which untraced records the original runs consumed.

Fairbench gives benchmark authors a rule for deciding what a diagnostic score may reflect (Online Methods, “The Fairbench protocol”): test each patient only with knowledge that was publicly available before the prioritization being evaluated. The patient’s own observations are the input, not the knowledge: they must have existed at that prioritization, but need not have been published. Moving knowledge into learned weights, preprocessing or implementation rules does not exempt it from this requirement. Conversely, later software can be used if the knowledge it uses satisfies Eq. (1). Fairbench thus states a necessary information-boundary requirement for claims of historical prioritization performance, not a certificate of clinical validity or benchmark quality.

These scores are not prospective diagnostic yields: the cases were published after their genetic causes were identified, reported phenotypes can include findings collected once the cause was known, and the variant-aware experiments use simulated exomes. Publication dates are a practical reference for the prioritization time, not a reconstruction of the original visit. Fairbench removes the temporal contamination; with the other contamination sources controlled as well — related cases in both the knowledge database and the test set among them — the benchmark becomes a fair estimate of historical prioritization performance.

An interpretable benchmark of a gene-prioritization system lets readers check both the score and the knowledge behind it. The check needs the run’s artifacts preserved: patient inputs, case dates, the exact knowledge-database releases, software, settings and outputs. A benchmark claiming a historical information boundary following the Fairbench protocol should release the records needed to verify it: inputs, manifests and code make the claim auditable, though auditability alone does not demonstrate compliance. Where the boundary cannot be established, authors should report each knowledge database’s release dates alongside the case dates, identify uncertified components, and refrain from the assessment-time claim.

The knowledge-database release used in testing a patient can substantially change benchmark accuracy, with patient inputs and ranking algorithms fixed. The audit tied that ranking knowledge to the very publications reporting the tested cases. Fair-bench makes the intended information boundary explicit; reporting a score with that boundary, and the evidence to check it, lets readers judge what diagnostic ability the benchmark actually establishes.

## 3 Online Methods

### 3.1 The Fairbench protocol

Fairbench ties the knowledge permitted in an evaluation to the prioritization being evaluated. The reference time is the time specified for the prioritization task represented by the case, independently of when the computational evaluation is performed or which resource release is scored. Let *p*(*t*) denote the patient information available at that prioritization, at time *t*. These observations need not have been published. Let *D*(*t*) denote all knowledge the diagnostic system uses to interpret *p*(*t*), and *K*(*t*) all knowledge publicly available strictly before *t*. Fairbench requires:

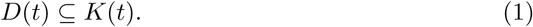

The requirement applies regardless of how knowledge is represented or supplied— for example, through databases, learned parameters or knowledge encoded in algorithms. Availability must be traceable through any preprocessing: transformations may not add information first publicly available at or after *t*. A later software implementation is admissible if all knowledge it uses satisfies Eq. (1).

Public release supplies the evidence for the cutoff. A qualifying preprint or authenticated resource release can establish availability; each database record need not have a separate publication citation. Our experiments use case publication as the reference because the dates of the original prioritizations are generally unavailable. The selected prepublication resources predate the cited case reports, but availability of every fixed component and patient observation has not been certified. The original date extraction also did not systematically search for earlier preprints or other reports of the same patient. These limits separate the resource comparisons performed here from complete compliance of a pipeline with Eq. (1).

### 3.2 Controlled comparison of dated knowledge resources

We compare rankings of the same published patients under prepublication and later knowledge-database states. We selected Exomiser, LIRICAL, Phrank and Phen2Disease for executable implementations and recoverable dated resources [15–18]. This was a practical selection, not a systematic screen of all available tools. The study cutoff for cases and resources was 23 October 2026. The last recovered releases predating it were dated 13 May 2026 for Exomiser and 3 September 2026 for the other systems.

We draw patients from public collections of published rare-disease cases, as described in Section 3.10. For each diagnostic system, eligible cases must have the required patient inputs, a reported causal gene and a verified publication date precise enough to select an unambiguous prepublication system data snapshot. We fix the eligible patient set, the patient inputs, the algorithm implementation and its configuration for each system. Only the supplied data snapshots change.

We also compare the systems on the intersection of their eligible patient sets, using each patient’s baseline and each system’s final available snapshot. To trace accuracy over successive releases, we select cases that can be compared at the system’s earliest later snapshot and keep that patient group fixed across all subsequent releases. These descriptive analyses were defined after the experiments were completed.

We use the same testing procedure for all four systems. Snapshot dates follow the release schedules of the data sources used by each system.

For each patient, we first run the system with the most recent knowledge-database snapshot released strictly before their case report. This run provides the patient’s prepublication baseline. We then test the same patient with every later available system data snapshot through the study cutoff. We record the rank of the reported disease-causing gene in each run.

At each later snapshot, we compare accuracy with baseline accuracy for the same patients. Because the patient inputs, algorithm implementation and configuration remain fixed, any accuracy difference is attributable to the resource intervention within this fixed pipeline. The contrast includes the combined effects of additions, corrections, removals and representation changes in those resources; it does not isolate every postpublication discovery.

### 3.3 Identifying database findings linked to the tested patients

We checked whether the knowledge databases contained phenotype associations linked to the publications describing the tested patients. We used the database files recorded in the original runs and verified their identities against saved file hashes. For Exomiser, we exported human disease profiles through its native model loader. Because these profiles do not retain citations, we matched their disease–phenotype associations to the source annotations identified by the database provider. The final phenotype database, release 2602, used HPO annotations dated 8 January 2026. For LIRICAL, Phrank and Phen2Disease, we examined the 3 September 2026 annotations and the mappings used to construct their disease or gene profiles.

We classified the recorded sources of each association before joining them to benchmark outcomes. For each patient, we examined phenotype associations across the diseases linked to the causal gene. We combined all recorded references for each association. An association was classified as exclusive to the patient’s publication only when every recorded reference resolved to that publication. Mixed, missing or unparseable references prevented this classification. A citation links an association to a publication, which may describe several patients; it does not identify which individual supplied each finding.

Table 1 includes patients with at least one such exclusively supported association. The audited cohorts contained 5,886 Exomiser, 1,090 LIRICAL, 3,062 Phrank and 3,071 Phen2Disease patients. Patients without a traced link remained in the full-cohort denominator. Failure to trace a link was not treated as evidence that the diagnostic knowledge was independent of the patient.

### 3.4 Checking how those findings enter scoring

We checked the role of the identified associations by replaying the systems’ native phenotype calculations. For Exomiser 15.1.0, we reconstructed the original patient inputs and replayed the human-phenotype scoring component used by the evaluated pipeline. For LIRICAL 2.4.1, we replayed phenotype likelihood calculations using both observed and explicitly absent patient findings. For Phrank and Phen2Disease, we verified that the audited associations reproduced the phenotype profiles supplied to their ranking implementations.

Complete diagnostic outputs provided a separate check on the component calculations. We compared the replayed calculations with retained outputs for selected patients and checked whether the causal gene appeared in the final ranking. We distinguished an association present in a database from a contribution to phenotype scoring and from confirmed use in a complete diagnostic output. A phenotype-score contribution alone does not establish a change in the final gene rank. The supplementary section “Publication sources and native scoring” (“Native phenotype scoring and full-output checks”) gives the validation scope and the accompanying LIRICAL example.

### 3.5 Accounting for accuracy in the identified patient groups

We joined the source classifications to each patient’s saved prepublication and latest results. Accuracy within an identified group was its summed fractional Top-1 credit divided by the number of patients in that group. Its contribution to the overall accuracy increase was the sum of its patients’ paired credit changes, divided by the full system cohort size and multiplied by 100. Contributions are therefore expressed in percentage points. Execution errors were never converted to zero credit.

We estimated 95% confidence intervals by resampling the full cohort’s source publications 10,000 times, keeping each publication’s patients, paired outcomes and group memberships together. Each replicate used its sampled full-cohort denominator; intervals were the 2.5th and 97.5th percentiles. Combined groups were calculated within each replicate rather than by adding interval endpoints. Supplementary Table S13 separately accounts for final credit in all four publication-support groups for Phrank and Phen2Disease. These post hoc analyses account for observed score changes without altering databases or rankings. They do not estimate how much of the accuracy increase was caused by findings from the tested patients’ publications.

### 3.6 Removing sole-source associations at the final knowledge-database release

The audit classifies each consumed phenotype association by its recorded sources. The removal turns that classification into an intervention: delete the associations supported only by the tested patient’s publication, and measure what Phrank loses.

Phrank is the system where this is verifiable, because its gene–phenotype resource names the supporting publications of every association. Exomiser’s consumed profiles carry no citations, and a LIRICAL deletion at a fixed release is already reported [13].

Patients, implementation, settings and the knowledge database stay exactly as in the main experiment’s final state (the HPO release of 3 September 2026). For each of the 1,690 case publications we deleted every association that publication alone supports — across all diseases and genes, not only the tested patient’s own — rebuilt Phrank’s model, and re-ranked that publication’s patients. Each patient is scored twice: with the full release, and with their publication’s sole-source associations removed. Edge cases — mixed, missing and negated source records, and deletions that leave the model unchanged — are defined in the supplementary section “Publication sources and native scoring” (“Source removal: the exact rule and what it changed”); the released plan preserves the exact rule.

Results are reported separately for the temporal experiment’s 3,062-patient Phrank cohort and for all 9,588 catalogued patients with an identified causal gene. A patient whose causal gene loses every association scores zero and stays in the denominator. With nothing removed, the sealed execution (23 November 2026) reproduces every cohort patient’s final-release result exactly, and an independent check confirmed that the deleted associations are exactly those the audit attributes to each publication alone. Intervals are 95% publication-cluster bootstrap intervals, as in the main analysis.

### 3.7 Baseline annotation status and between-system comparisons

A patient counts as unannotated at baseline when the original annotation files of their baseline release hold no phenotype association for the causal gene. We read this from the files themselves — the same files the run used, verified by checksum. For Exomiser we read only the human phenotype data of its native profile, because its animal-model and interaction evidence can rank a gene without any human annotation. A system’s configuration can also read less than the files hold — LIRICAL’s does not read Orphanet-linked entries — so the supplement reports, beside the file-based status, what each system actually saw. Group accuracies and contributions carry the same 95% publication-cluster confidence intervals as every other analysis (“Accuracy measures and statistical analysis”). The between-system comparison uses the shared cohort, where the same patients run in all four systems: accuracy differences are paired patient by patient, at the historical baseline and at each system’s final release.

### 3.8 Reassessment of published benchmarks

We examine how accuracy changes when cases from published benchmarks are retested with knowledge databases released before their case reports. We use the original patient cases and public materials from the Exomiser evaluation in PhEval and from studies of LIRICAL, Phen2Gene and Phen2Disease [14–16, 23].

We first attempt to reproduce the authors’ results using the original patient inputs, algorithm implementation, configuration and data. Where original resources cannot be recovered, we reconstruct the reference and document the differences. We distinguish comparisons against a reproduced benchmark from comparisons against a reconstructed reference.

For the Exomiser reference, we reproduce the PhEval phenotype-only preset using Exomiser 14.0.2 and release 2406 resources. Gene scores are rounded to four decimal places before descending worst-tie ranks are compared. Agreement refers to this precision and convention; the main experiment instead uses fractional tie credit. Detailed reference comparisons and the excluded mismatch are included in the released verification records.

We then retest eligible cases with recoverable data snapshots released strictly before each patient’s case report, keeping the patient inputs, algorithm implementation and configuration fixed. Cases require verified publication dates and dated earlier snapshots. Eligibility is determined before the prepublication arm is measured. Phen2Gene is excluded from the reported temporal comparisons because its rebuilt reference did not reproduce the released term tables and weights.

Both conditions use the same patients and scoring rules. Results report the included cases and measured changes; Supplementary Notes document reconstruction limits and case-level exclusion decisions.

### 3.9 Accuracy measures and statistical analysis

We measure changes in ranking accuracy on the evaluated cases. Accuracy inflation refers to an overestimate relative to a specified information boundary; the experimental contrasts below directly measure differences between recovered resource states.

*Our main experiment.* Top-1 causal-gene ranking accuracy is the primary measure; Top-3 and Top-10 are supporting measures. For each system, we compare each later resource snapshot with the prepublication baselines of the same patients.

To avoid rewarding a system for leaving many genes tied at the top, we average success over all possible orderings within each group of tied genes. When a group crosses the Top-*k* cutoff, each gene in it receives the same partial credit. Thus, ten genes tied first give the causal gene 0.1, 0.3 and 1.0 credit at Top-1, Top-3 and Top-10, respectively. The same rule applies to baseline and later snapshots.

*Reassessment of the authors’ benchmarks.* Each pair uses the same patients, ranked target, thresholds and scoring rule. We retain the original endpoint where verified: genes for Exomiser and diseases for LIRICAL. Phen2Disease uses a reconstructed gene-ranking endpoint with the same fractional tie rule in both arms because the original tie convention was not recovered. Supplementary Notes describe the references and reconstruction limits.

For the temporal comparisons, we subtract accuracy with prepublication data from accuracy with later data, using the same cases. We express the difference in percentage points.

If the reported causal gene or diagnosis is absent from the candidate list, the case counts as a failure. Execution errors are not scored as diagnostic failures and prevent a run from being finalized.

Exploratory 95% confidence intervals use 10,000 bootstrap replicates. We resample source publications, keeping all cases from each sampled publication together and retaining each patient’s paired results. Each replicate estimates the patient-weighted mean accuracy difference; intervals use the 2.5th and 97.5th percentiles. Intervals are not adjusted for multiple comparisons and do not include uncertainty in publication dates or resource reconstruction. Seeds and calculation details are provided in Supplementary Methods, “Accuracy, decomposition and uncertainty.” Supplementary Table S10 reports a separate sensitivity analysis clustering the variant-aware results by background donor. The analysis plan was exploratory: analyses were defined after the initial experiments, an execution manifest was frozen before the clean reproduction run, and no preregistration exists.

### 3.10 Patient cohort for the main experiment

To compare resource effects across a broad range of rare diseases, we assembled a common patient collection for the main experiment. We obtained 10,377 case records from Phenopacket Store [19]. These records link patient phenotypes, represented using the Human Phenotype Ontology (HPO) [20], and reported diagnoses to published case reports. Each system evaluates only cases with an available data snapshot released strictly before the case report. The study analyses previously published, de-identified case reports and public databases; no new human-participant research was conducted. We use the earliest documented publication date of the cited case report as the reference date *t*, because diagnosis dates are generally unavailable. Dates are obtained from PubMed and, where needed, publisher records. The date-invariance rule for uncertain dates was applied after inspection of the original outcomes; adjudication used source evidence, not ranking performance. Two dating channels limit what the compared states can contain. A prepublication release can omit knowledge already published before the case report, and a publication-dated baseline can include knowledge unavailable at the original assessment. Neither fact, on its own, establishes the direction of the resulting accuracy difference. Date-uncertain cases were excluded unless every plausible date selected the same snapshot; conversion and reconstruction residuals likewise have no established direction and remain uncertified.

When only the publication month is known, we retain the full month as the possible publication period. We exclude cases whose publication dates cannot be verified or are too imprecise to select a single prepublication snapshot. Publication dates were verified, and corrected where needed, before eligibility and baseline selection for the definitive analysis; cases whose unresolved dates could change eligibility or baseline selection were excluded from the affected system’s estimates.

Historical coverage determined how many patients each system could evaluate. Each patient’s baseline used the most recent usable release strictly before their report; cases without such a release were excluded from that system’s comparison. The recoverable releases span different periods, yielding different eligible cohorts. Table 3 gives the earliest usable release and the number retained after publication-date and patient-input checks.

**Table 3.** Patients included in each main experiment from the common catalogue of 10,377 cases. Counts reflect snapshot availability, publication-date verification and required patient inputs.

| System | Earliest usable dataset snapshot | Patients included |
|---|---|---|
| Exomiser | 12 December 2017 | 5,886 |
| LIRICAL | 1 July 2024 | 1,090 |
| Phrank | 5 October 2022 | 3,062 |
| Phen2Disease | 5 October 2022 | 3,071 |

To compare resource effects on the same patients across systems, we also evaluated the 1,090 cases eligible for all four systems. This is exactly the LIRICAL-eligible cohort: 108 publications and case reports after its earliest usable snapshot of 1 July 2024. The recoverable compatible LIRICAL resource series starts on that date. Shared membership controls which patients are compared, but does not equalize resource dates or make the cohort representative of all rare-disease cases.

LIRICAL uses both observed and explicitly absent phenotypes at every release. One patient with conflicting phenotype signs at baseline was excluded from its paired comparison; execution failures were not counted as diagnostic failures (Supplementary Methods).

For systems requiring variant input, we construct simulated exomes by combining each case’s reported causal variants with a verified background VCF. Backgrounds are derived from the OpenScientist–Exomiser benchmark archive [24]. Source versions and preparation details are given in Supplementary Methods.

### 3.11 Knowledge resources and snapshot selection

To apply the temporal restriction in Eq. (1), we recover historical releases of the data used by each diagnostic system. We collect these snapshots from public archives up to the study cutoff and verify when their required components became available.

Exomiser’s phenotype, interaction and genome resources vary across snapshots. LIRICAL varies its HPO annotations, gene–disease mappings and variant resources, but uses fixed engine transcripts and bundled background frequencies. Phrank and Phen2Disease vary HPO ontology and gene–phenotype resources. Identifier mappings and algorithm implementations are held fixed. The historical availability of every embedded parameter and auxiliary resource has not been certified. Supplementary Table S1 summarizes this scope.

Older formats sometimes require conversion for the fixed implementation. Source and output identities and conversion checks are retained. A recent conversion date alone does not imply that later diagnostic knowledge was introduced; conversely, an old source does not certify every transformation. Supplementary Methods describe the recovered releases and remaining provenance gaps.

### 3.12 Established-association sensitivity

To assess heterogeneity within the temporal results, we compared cases with independent curated evidence of an established association against those classified from an earlier report in the collected source cases. These original evidence-basis labels are not a complete history of gene discovery or a clinical population classification. They do not determine gene ranks or eligibility for the overall temporal comparison. Supplementary Table S12 reports the outcomes for these groups.

### 3.13 Use of large language models

Large-language-model agents, directed and reviewed by the author, contributed substantially to the code and its review, the analyses and the drafting of this work. The agents ran on Claude Fable 5 (Anthropic) and GPT-6-Astra (OpenAI), used between September and November 2026, each model cross-checking the other’s output. The author conceived and directed the study, set the data-adjudication rules, verified the results and takes responsibility for the work; interpretation and conclusions remained with the author. No reported value was produced by a language model: every number derives from the released pipeline and is covered by its verification records.

## Data availability

Patient records are available from Phenopacket Store release 0.1.27 and background exomes from the OpenScientist–Exomiser archive, version 2 [19, 24]. Case-level results, date evidence, selection decisions, resource identities, figure source data and the verification records of the reported run are archived at Zenodo (https://doi.org/10.5281/zenodo.24503734), together with an archive of the code repository. Third-party resources are linked to their original archives; files excluded for licensing are listed in the deposit with the reason for each.

## Code availability

The evaluation harness, historical-resource converters, analysis code and container configurations are available at https://github.com/matevz-kovacic/fairbench and archived in the deposit above. The release identifies the source revision and container images of the reported run and documents the command sequence that rebuilds the images and regenerates the study from public inputs. The removal experiment is part of the same release, with its exact rule, code and verified outputs, and so are the follow-up analyses.

## Funding

Resources for this work were provided by Inetis Ltd.

## Author contributions

M.K. conceived and designed the study, directed and verified all analyses, and wrote the manuscript.

## Ethics

This study analysed previously published, de-identified case reports from a public corpus; no new data from human participants were collected and no ethics approval was required.

## Competing interests

The author is employed by and holds an ownership interest in Inetis Ltd., which provided resources for this work. The author also develops a separate gene-prioritization method, not evaluated here, which could benefit from changes to benchmarking practice proposed in this work.

## Extended Data

**Extended Data Fig. 1.**
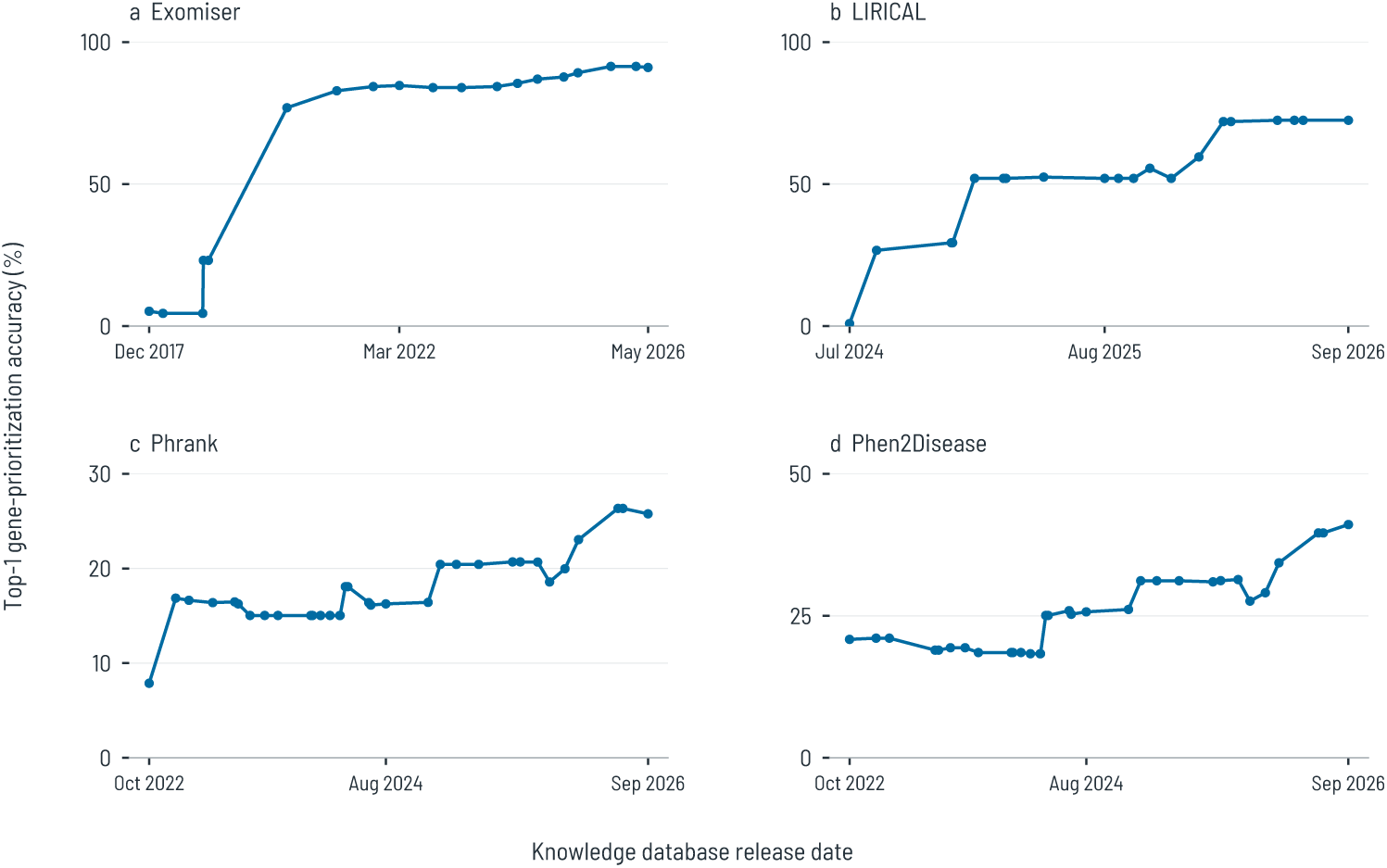
Illustrative knowledge-database release trajectories. Fixed groups contain 268 Exomiser cases, 225 LIRICAL cases and 475 cases each for Phrank and Phen2Disease. Curves begin at their shared prepublication resource baseline. These groups are concentrated: 157 NF1 cases from one publication constitute 58.6% of the Exomiser panel; three publications supply 70.7% of the LIRICAL panel. Each Phrank and Phen2Disease panel contains 256 ANKRD11 cases from one publication (53.9%); its three largest publications supply 67.2%. The groups are illustrative, not representative clinical samples. Axis scales differ. The supplementary section “Publication sources and native scoring” (“NF1 and the Exomiser release components”) examines the concentrated Exomiser trajectory.

**Extended Data Table 1.** Phrank accuracy at the fixed final release before and after removal of the associations supported only by each patient’s publication. Top-*k* accuracy in percent; decreases in percentage points with 95% publication-cluster confidence intervals (10,000 draws), calculated before rounding. All accepted patients: every catalogued patient with an NCBI Gene identifier for the causal gene (9,588 patients, 1,690 publications). Matched temporal cohort: the patients with a final-state result in the temporal experiment’s Phrank arm (3,062 patients, 351 publications); the two populations are reported separately throughout. Patients whose causal gene loses every association after the removal, or has none at the final release, remain in the denominators with zero credit (Online Methods).

|  | Untreated final (%) | After removal (%) | Decrease, pp (95% CI) |
|---|---|---|---|
| *All accepted patients (9,588 patients; 1,690 publications)* |  |  |  |
| Top-1 | 52.51 | 23.42 | 29.09 (24.89–33.43) |
| Top-3 | 63.70 | 33.93 | 29.77 (25.33–34.32) |
| Top-10 | 74.35 | 45.92 | 28.43 (24.02–33.02) |
| *Matched temporal cohort (3,062 patients; 351 publications)* |  |  |  |
| Top-1 | 60.81 | 9.97 | 50.83 (40.80–60.44) |
| Top-3 | 70.50 | 15.35 | 55.15 (44.29–65.66) |
| Top-10 | 79.22 | 23.51 | 55.71 (44.46–66.59) |

## Notes

https://doi.org/10.5281/zenodo.24503734

https://github.com/matevz-kovacic/fairbench

