## Supplement for "Standard benchmarking inflates gene-prioritization accuracy estimates"

### Fairbench: Supplementary Information

23 November 2026

This supplement specifies what the manuscript holds fixed and what it varies: the knowledge-database releases and inputs behind each comparison, the reconstruction decisions behind the reassessments, the uncertainty machinery, the sensitivity analyses, and the source audit with its native-scoring checks. Tables and figures use the corrected temporal cohorts; reassessments retain their separately verified patient collections. Confidence intervals for the main comparisons appear alongside their estimates in the manuscript's main results figure and tables. Throughout, "corrected" names the definitive cohorts and outcomes of the completed study; the term matches the released analysis records.

#### Supporting figures and tables

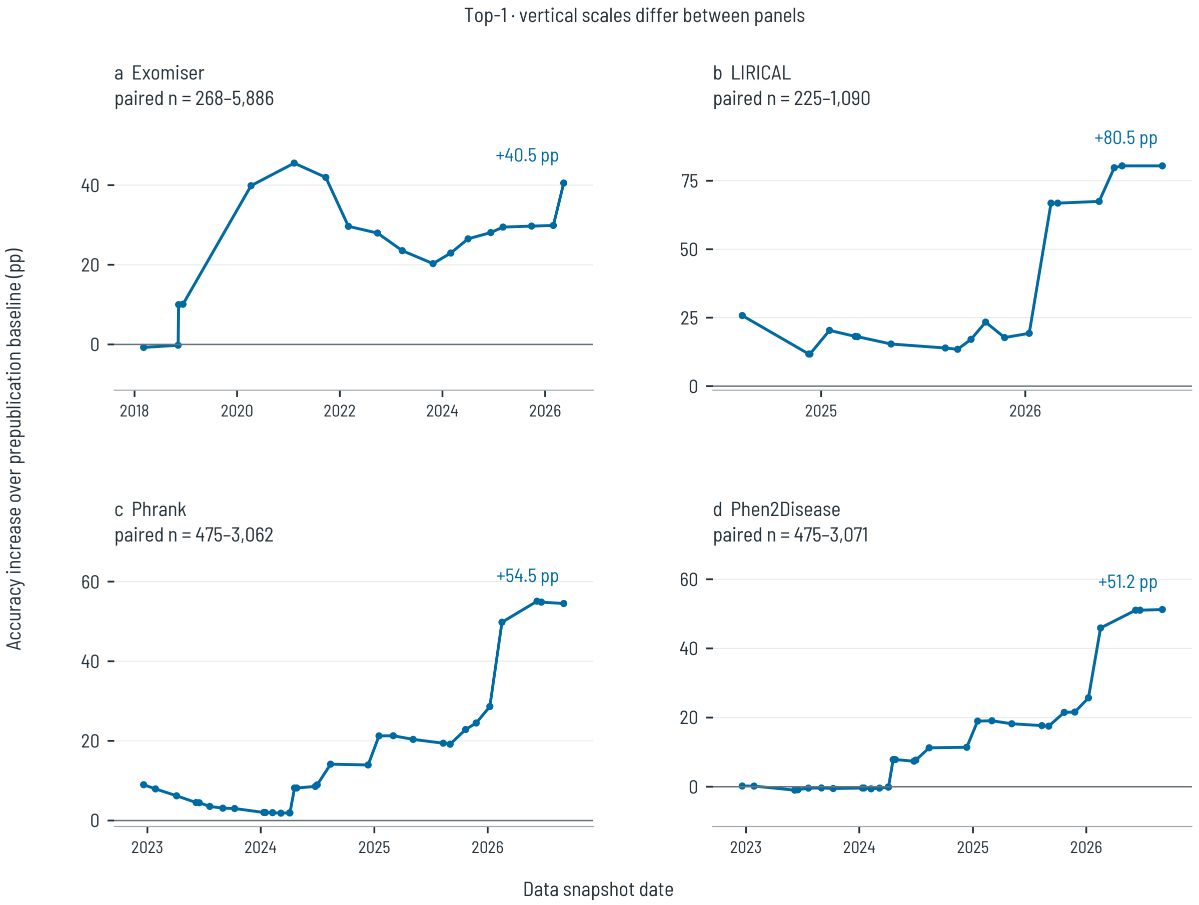


**Figure S1: Top-1 effects using all available paired cases.** At every point, the later results are compared with the baselines of exactly those patients. The number of patients grows as additional case reports fall before the snapshot date; the title gives its range. Changes between adjacent points can therefore reflect both updated data and changed patient membership. These curves complement the fixed-group trajectories in manuscript Extended Data Fig. 1. Vertical scales differ between panels. Lines connect measured releases without interpolation of patient results. Source data: `source-data/main-trajectories.csv`.

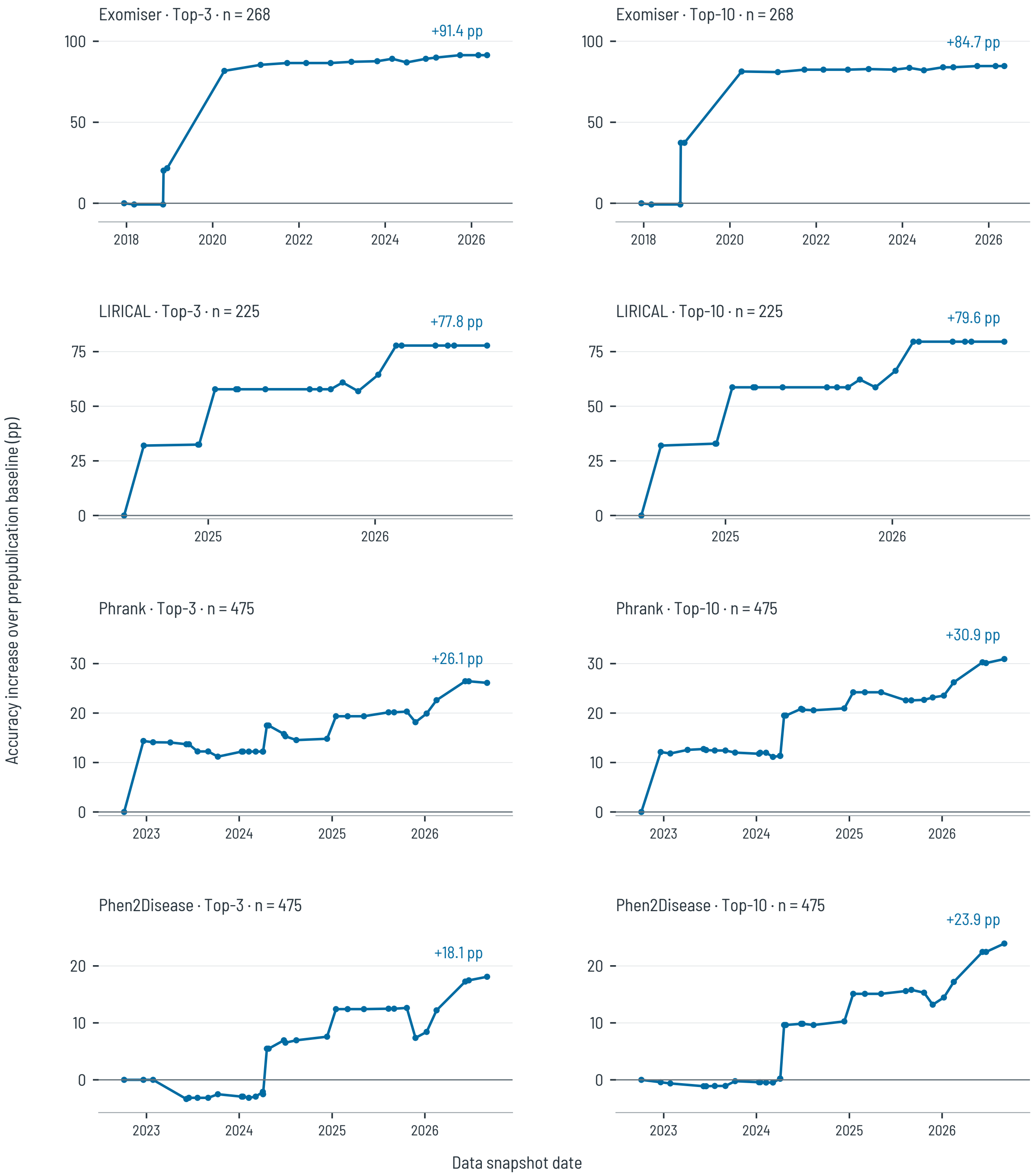


**Figure S2: Top-3 and Top-10 changes within fixed patient groups.** The patients and data releases are the same as in manuscript Extended Data Fig. 1. Each panel shows one endpoint to keep the trajectories distinct. The two panels within each system use the same vertical scale; scales differ between systems. Labels give the final increase over prepublication accuracy. Source data: `source-data/main-fixed-cohort-trajectories.csv`.

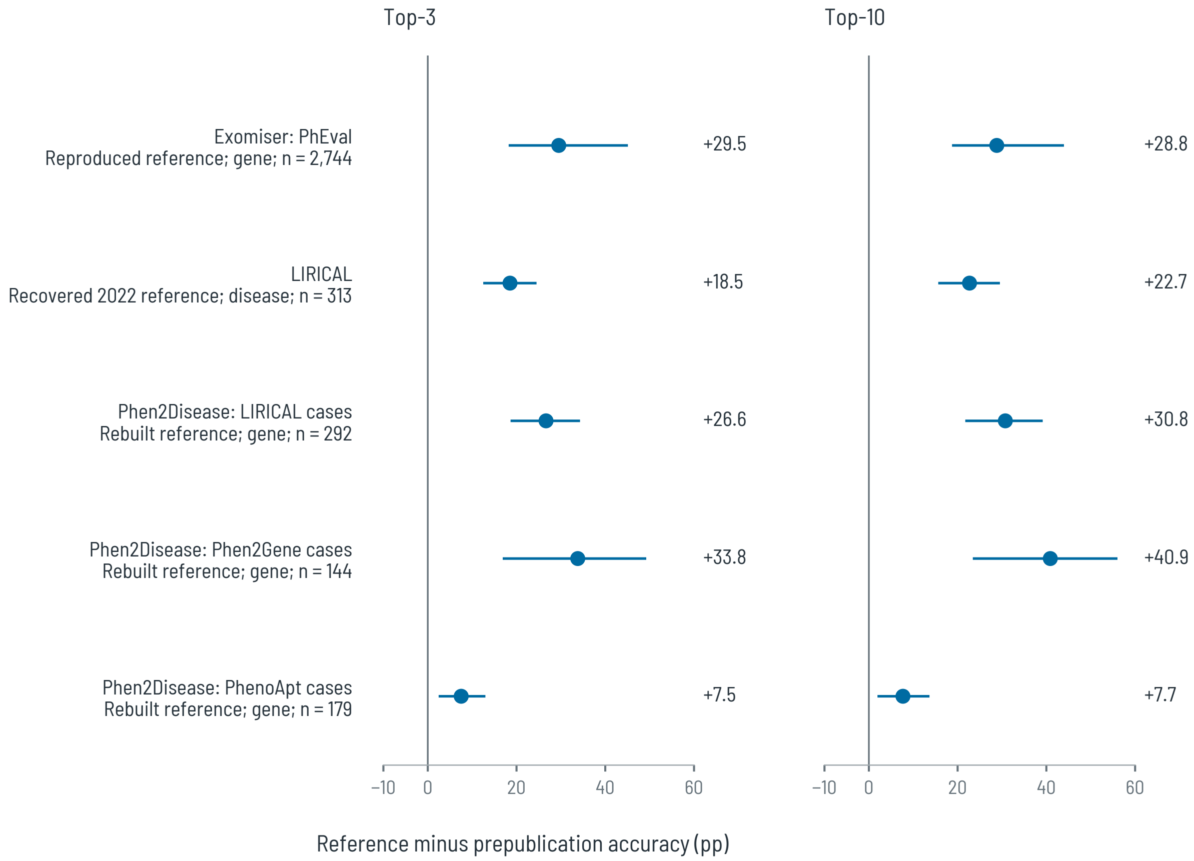


**Figure S3: Top-3 and Top-10 reassessment differences.** Patients and references match Table 2. Phen2Disease row labels name the source patient collections. Points show reference minus prepublication accuracy; lines show exploratory 95% publication-cluster bootstrap intervals. The panels share a common scale and show each endpoint separately. Positive values indicate greater accuracy with the later reference. Reconstruction and dating uncertainty are not included in the intervals. Source data: `source-data/reassessment-summary.csv`.

### Supplementary Notes

#### Scope of historical knowledge-database comparisons

The experiments hold patient inputs, software and configuration fixed while replacing specified knowledge databases. Table S1 distinguishes these interventions from full certification of the information used by the pipeline. It summarizes the code-facing inventory in `paper/reviewer-followup/resource-scope.csv`; release identifiers, checksums and conversion receipts remain in the completed-run records.

**Table S1:** Varied resources and fixed components in the main study. Archive dating establishes the availability of the selected releases; it does not identify every publication underlying their records. Fixed means unchanged between the paired runs, not historically certified.

| Scope | Resources varied | Fixed components or unresolved scope |
|---|---|---|
| Exomiser | Phenotype and model-organism models, matched interaction resource, genome variant and ClinVar stores, genome transcripts | Implementation and configuration; publication-level attribution in aggregated variant and phenotype resources is incomplete. |
| LIRICAL | HPO ontology and annotations, gene–disease mappings, variant and ClinVar stores | Engine transcript resources and bundled background frequencies are not historically certified for every patient cutoff. |
| Phrank and Phen2Disease | HPO ontology and gene–phenotype annotations | Fixed implementations; compatibility of patient terms with historical ontologies requires separate assessment. |
| All systems | The resources listed above | Case-report inputs and reported diagnoses, identifier mapping, implementation and configuration remain fixed. Historical provenance of every embedded parameter has not been established. |
| Variant inputs | No change within a patient pair | Reported causal variants combined with benchmark background VCFs; these are simulated exomes, not reconstructed historical clinical sequencing records. |

The fixed HGNC identifier resource is pinned to 6 October 2026. A modern identifier representation is not itself proof of diagnostic leakage, but information-bearing changes in mapping require an audit. Likewise, conversion for a modern implementation must preserve the intended source information. Retained source identities and native conversion checks support reproducibility without certifying every unresolved route as historically valid.

#### Published-benchmark reconstruction

**Why some cases could not be retested.** A temporal comparison requires a verified publication date and recoverable knowledge-database releases from before that date. An imprecise date is acceptable only if every possible date selects the same earlier snapshot. Other exclusions concern missing or conflicting patient inputs, unresolved gene or patient identities, duplicate records, and an Exomiser case whose original result was not reproduced. The recorded decisions cover every source entry, including exclusions, in the source-data files listed below. Counts refer to entries in each benchmark and are not pooled across overlapping patient collections.

- Exomiser: `reassessment-exomiser-eligibility.csv`.

- LIRICAL: `reassessment-lirical-case-decisions.csv`.
- Phen2Disease: `reassessment-phen2disease-case-decisions.csv`.
- Phen2Gene (excluded algorithm): `reassessment-phen2gene-case-decisions.csv`.

**Exomiser.** The reproduced reference is PhEval's phenotype-only Exomiser arm: Exomiser 14.0.2, the native `phenotype-only` preset and release 2406 resources. Scores were rounded to four decimal places before descending worst-tie ranking. Thus, agreement concerns the deposited precision and tie convention, not unrounded floating-point scores. This differs from the variant-aware main experiment.

Scores and ranks matched for 4,676 of 4,677 cases. The exception, `PMID_17886299_patientBM5`, had deposited rank 18 and replay rank 16, with candidate counts 16,065 and 16,255. Its cause remains unresolved, and the case is excluded from the historical comparison. The matched full-reference Top-1 accuracy was 39.09% (1,828/4,676).

Verified dates and recoverable historical resources retained 2,744 cases from 213 publications. Their reference Top-1 accuracy was 28.43% (780/2,744), versus 3.57% (98/2,744) historically. The 24.85-point difference has a 95% publication-cluster interval of 15.90–38.85 points. There were 699 gains and 17 losses. Top-3 and Top-10 differences were 29.52 and 28.83 points. These values apply to this retained phenotype-only comparison; they do not estimate variant-information contributions by comparison with the main cohort.

The archive includes paired outcomes, 4,677 detailed reproduction records, mismatch details, archived execution code and a verifier in `paper/reviewer-followup/exomiser-reassessment-evidence/`. A separate recalculation implementation reproduces the estimates and the original 10,000-draw bootstrap.

**LIRICAL.** For LIRICAL, 313 of 384 public cases could be paired. Using the original native disease-ranking endpoint, Top-1 accuracy was 22.0% with the recovered 2022 reference and 10.5% with prepublication data, a difference of 11.5 percentage points (95% interval, 5.7–17.4). Top-3 and Top-10 differences were 18.5 and 22.7 percentage points (Fig. S3). The original 2019 annotations were not recovered, so these estimates describe the effect on the original cases under the recovered reference rather than a correction to the paper's reported accuracy.

**Why Phen2Gene was excluded.** The 281 released Phen2Gene patient outputs were reproduced, but regenerating its term tables and weights did not reproduce the reference. We therefore exclude the algorithm from temporal-effect estimates. The attempted reconstruction and case decisions remain archived. This exclusion does not concern Phen2Disease's use of the Phen2Gene patient collection.

**Phen2Disease.** Phen2Disease's three assessable patient collections came from the LIRICAL public literature cases (384 entries), the Phen2Gene benchmark (281 entries), and PhenoApt source cohort 1 (185 entries). The workflow identifies these as cohort1, cohort2 and cohort3, respectively. These identifiers denote patient collections, not database releases. The input archives and their checksums are recorded in the frozen resource inventory.

Historical comparisons were possible for 615 of 2,051 source entries across the six original collections: 292 LIRICAL cases, 144 Phen2Gene cases and 179 PhenoApt cases. Top-1 accuracy fell from 37.2% to 12.7% on the LIRICAL cases and from 30.9% to 3.5% on the Phen2Gene cases, decreases of 24.6 and 27.4 percentage points. On the PhenoApt cases, accuracy changed from 23.1% to

21.0%, a difference of 2.1 percentage points whose 95% interval included zero. Top-10 differences were 30.8, 40.9 and 7.7 percentage points, respectively.

The original similarity matrix and evaluation tie convention were not recovered. These are paired reconstructions using the same fractional tie rule in both conditions, not exact corrections to the published results. Patient collections were analysed separately because source cases may overlap.

**Scope of the comparisons.** Across these experiments, later database states changed ranking accuracy while patient inputs and the selected algorithm were held fixed. The comparisons measure the effect of the recovered database states. They do not certify the historical provenance of every fixed software component, or distinguish all later additions from database corrections and removals. The complete inclusion decisions, reconstruction limits and numerical results accompany the tables and supporting figures.

#### Clinical applicability and historical interpretation

**Patient selection and inputs.** We used published, solved cases and simulated exomes. These need not represent the patient mix, phenotype documentation or sequencing data encountered in a particular clinic. Reported phenotypes may include findings obtained during diagnosis; we did not establish when every patient feature became available. The study therefore does not replace prospective evaluation in a representative clinical cohort.

**Publication dates as practical references.** Case-report publication dates were used because the dates of the prioritizations being evaluated were generally unavailable. A report may appear after that prioritization. Consequently, a database released before the report could include knowledge unavailable during the actual assessment. This limits how closely our experimental cutoff represents the original date; it does not change the Fairbench requirement to respect the time of the prioritization being evaluated.

**Archive coverage and fixed resources.** The recoverable releases cover different periods for each system, so shared patients can still have different baseline dates. A released snapshot can omit knowledge already published, and later releases can correct or remove records as well as add discoveries. The measured accuracy difference is the combined effect of replacing the recovered database states. We did not verify the historical availability of every resource or parameter held fixed. The observed increases therefore cannot be assumed to be minimum estimates of temporal contamination, or interpreted as a ranking of the systems' clinical usefulness.

### Supplementary Methods

#### Patient sources, inputs and eligibility

The common input collection contains 10,377 records from Phenopacket Store release 0.1.27. Its archived release is pinned by a SHA-256 checksum in the source inventory. Patient identifiers connect the records to their cited publications, phenotypes and reported causal genes. They identify source records; they do not by themselves prove that individuals described in different publications are distinct.

Simulated exomes combine the reported causal variants with background VCFs obtained from the separate OpenScientist–Exomiser benchmark archive, Zenodo record 22967674, version 2 (`https://doi.org/10.5281/zenodo.22967674`). Background cases are not added to the phenopacket cohort. The same prepared patient inputs are used at both endpoints of a comparison. Causal genes are used as evaluation targets; adding a reported causal variant to an exome is part of the simulated-input design, not evidence of an independently ascertained clinical exome.

Each system starts from this common collection. Eligibility requires a preceding usable knowledge-database state, a sufficiently precise verified case-report date and inputs accepted by the preparation workflow. The resulting cohorts contain 5,886 Exomiser, 1,090 LIRICAL, 3,062 Phrank and 3,071 Phen2Disease cases; their intersection contains 1,090 cases. Recorded exclusions include shared input-preparation rejections as well as system-specific timeline restrictions. A variant-related reason in the shared preparation record does not mean that every algorithm consumes variant input.

The exclusion files assign a recorded reason according to the workflow's checking order. These counts are not independent counts of every defect a case might have. Full main-study reason counts are supplied in `source-data/main-exclusions.csv`, with additional correction exclusions in `source-data/corrected-exclusions.csv`; case-level reassessment decisions are listed in "Published-benchmark reconstruction." Cases not returned by a successfully executed ranking remain in the denominator.

#### Patient dates and baseline selection

The completed study used the earliest documented publication date of each cited case report, supported by PubMed and publisher records. A partial date is represented as a bounded interval. A case is eligible only when all dates in that interval select the same strictly preceding snapshot; a same-day release is ineligible. Diagnosis dates are generally unavailable, and the completed workflow did not exhaustively search for earlier reports of the same patient.

Publicly accessible preprints qualify as prior knowledge. The cutoff concerns when information was available, not whether a later publication was the first to establish a gene's causal role.

An earlier report about other patients does not move the evaluated patient's reference date. A date correction requires evidence that the earlier report describes the same individual. Findings added in later case descriptions remain an input limitation. Earlier cutoffs can change snapshot selection or eligibility; their effect on accuracy is not assumed monotonic.

The patient-date adjudication, including PGBD5 and the RNU5B-1/RNU2-2 reports, is recorded in `date-adjudications.csv` under the submission-closure archive. Changed cutoffs were propagated into baseline selection and native execution; unresolved comparisons were excluded.

The definitive temporal estimates retain an uncertain-date case only when all evidence-supported dates preserve eligibility and select the same baseline for that system. Comparisons whose dates selected different baselines, altered eligibility, or could not be bounded sufficiently to establish invariance were excluded from the definitive primary estimate, pending resolution; evaluable alternatives were reported separately. Cases ineligible under every plausible date followed the ordinary exclusion rule. Recency restrictions, elapsed-time coordinates and shared-cohort membership require their own invariance checks. Each paired temporal estimate uses the same retained patients in both conditions. This rule was applied after inspection of the original outcomes; adjudication used source evidence, not ranking performance. It addresses the identified disputed dates, rather than certifying every patient's first report.

Of the 5,886 distinct patients of the four cohorts (823 case publications), 4,361 patients' case-report dates (669 publications) come automatically from the frozen PubMed records, as the earlier of the print and electronic date. The remaining 1,525 patients' dates were set or verified from publisher or source evidence: 1,511 (151 publications) against archived publisher date pages, and 14 (3 publications) from verified earlier reports of the same patients. Rather than a random sample, every automatically assigned date was re-derived from the frozen PubMed records with the pipeline's own rule and matched the catalogue in all 4,361 cases; all 151 publisher-evidence dates match their archived evidence. The accounting is recorded in `paper/reviewer-followup/date-audit-counts-20261123.json`.

#### Implementations and reproducible execution

The frozen main-study records identify Exomiser 15.1.0, LIRICAL 2.4.1, Phrank 1.6 and Phen2Disease source commit `2c117fba11f059ce8e4a56de2ac296e3fba19405`. Implementations, patient inputs and analysis configurations are fixed within each paired comparison. The study cutoff is 23 October 2026; the last recovered releases predate that cutoff, as stated in the main tables.

Exomiser uses the pinned exome-analysis preset. LIRICAL's recorded configuration specifies REFSEQ transcripts, lenient validation and non-global analysis. Lenient validation runs LIRICAL's own sanitizer against each historical ontology, retaining its warnings — for example, terms unknown to an older release — while native input-validation errors make a case ineligible; execution failures are handled separately and are not entered as diagnostic failures. Non-global candidate selection and REFSEQ transcripts follow the released default configuration. Phrank uses direct gene ranking, with candidates from snapshot-annotated genes. Phen2Disease uses the released integrated scoring blocks and its declared strict-ancestor Lin convention, with candidates derived from the same gene–phenotype rows used to construct disease profiles. LIRICAL's main workflow uses both HPO terms and simulated VCFs; it is not phenotype-only. The retained manifests include implementation and input hashes and the detailed settings; these summaries do not substitute for them.

In the completed study, exploratory analyses preceded a frozen execution manifest; the definitive run validates saved cells against that manifest. Completed cells are retained with their candidate and tie evidence. Execution failures are not entered as diagnostic failures. Original outputs remain unchanged during the follow-up analyses. The downloaded results bundle and its inventory provide the input identities for the local analysis scripts.

The definitive run executed 19–23 November 2026 as one sealed, containerized launch: twenty-four jobs, each run once, every input fetched from its public provider and checked against a pinned or recorded checksum before use.

File paths in this document are those of the study working tree; every file cited is included in the data deposit, among the reported run's verified outputs under `main-experiment/` or under `records/`; the deposit inventory maps each printed path to its file.

LIRICAL uses observed and explicitly absent HPO terms throughout the corrected temporal comparison. Of its 1,090 included patients, 1,026 have explicitly absent terms, comprising 25,280 such terms in total. One patient with conflicting signs at baseline is excluded rather than assigned zero accuracy. All 17,763 included patient–release cells have valid ranked or not-ranked outcomes. The earlier runs that omitted absent terms are superseded for this arm. The separate published-benchmark LIRICAL reconstruction already preserved negative phenotypes and its estimates are unchanged.

#### Knowledge-database conversion and temporal scope

Table S1 describes the intervention. Archived sources are identified by release, source location, availability evidence and checksums. Compatibility conversions adapt historical representations to the fixed implementation. Recorded source and output hashes identify the converted artifacts. Native checks and conversion receipts are retained for inspection.

The comparisons do not certify the historical availability of every fixed auxiliary resource, fitted parameter or patient phenotype. Nor does a successful format conversion establish that all intended source information was preserved. These are distinct checks: format compatibility, information preservation and historical eligibility must not be treated as interchangeable.

All 32 identified Exomiser conversion checks passed their recorded preservation criteria. The interpretation against the captured analysis configuration found no discarded source selected for scoring. The per-release matrix and verification records are retained under `conversion-interpretation/` in the submission-closure archive. This validates the checked transformations and configured fields; it does not establish the historical availability of every fixed component.

#### Accuracy, decomposition and uncertainty

For a returned causal gene with $h$ genes scored strictly above it and $m$ genes sharing its score, Top-$k$ credit is

$$c_k = \min\{1, \max\{0, (k - h)/m\}\}.$$

An absent causal gene receives zero credit. Accuracy is the mean credit multiplied by 100. For a paired set of $n$ cases, the primary contrast is $100n^{-1} \sum_i (c_{i,\text{later}} - c_{i,\text{base}})$. This is a patient-weighted difference on the evaluated cases. Reassessment endpoints follow their stated reproduction or reconstruction rules; LIRICAL's reassessment ranks diseases.

The corrected main, recency, phenotype-compatibility and identity sensitivity intervals use 10,000 paired publication-cluster draws with base seed 20261109. Each system's seed label comprises its name and sorted retained patient identifiers, separated by newlines. The Exomiser reassessment uses the same base seed with the label `exomiser-reassessment`. These analyses add the integer represented by the first eight SHA-256 bytes of the label, in big-endian order, and use PCG64. The separately archived LIRICAL and Phen2Disease reassessments retain their original bootstrap settings. Source publications are sampled with replacement; all their cases and both endpoints remain together. Each draw computes the ratio of the summed paired differences to the sampled number of cases. The interval uses linear-interpolated 2.5th and 97.5th percentiles. No between-publication interval is estimated from a single cluster.

The established-evidence sensitivity uses base seed 20261111 with its recorded subset labels. Donor-cluster intervals use the main bootstrap implementation with background donors as the resampling unit and a separate system-specific label. All intervals are exploratory: they are not adjusted for multiple comparisons and do not incorporate uncertainty in dates, reconstruction or cross-publication patient identity. Alternative weighting summaries are descriptive means without new confidence intervals or clinical population weights.

The scripts and input hashes are recorded in `source-data/corrected-integration-verification.json`; reassessment settings are retained in `source-data/analysis-provenance.json`. Main estimates and intervals were recomputed from the corrected paired outcomes during manuscript integration. Reassessment outputs reproduced unchanged from their own hash-verified archive.

The publication-support groups let us account for both final accuracy and the temporal increase using the same patients. Their intervals use the same 10,000 publication-cluster bootstrap draws (PCG64, seed 20261111; publications ordered lexicographically within each system). In each draw, a contribution is divided by the sampled patient count, whereas a share of final credit is divided by the sampled total final credit. A missing group contributes zero, and no draw had zero total final credit. The intervals describe sampling variation with the recorded support held fixed. Input and native profile hashes and draw identities are retained in the verification record; the audit code defines the seed and publication ordering.

#### Supporting analyses of temporal comparisons

All results in this section use the corrected cohorts and saved outcomes, including the completed replacement runs. The supporting calculations require no further diagnostic execution. Their input hashes, reproduced main intervals and unchanged reassessment checks are recorded in `source-data/corrected-integration-verification.json`.

##### Release dependence and cohort concentration

Keeping the January already-evaluated cases fixed separates release effects from changing endpoint eligibility (Table S2). All three systems show a large February increase, followed by smaller changes. This table does not attribute each increase to an individual annotation.

**Table S2:** Top-1 increases over patient-specific historical baselines at three endpoints, holding January's cases fixed. Values are percentage points. These sets include cases already tested with resources later than their own baselines in January; January baseline cases are excluded.

| System | Cases | 8 Jan | 16 Feb | 3 Sep |
|---|---|---|---|---|
| LIRICAL | 1,063 | 19.29 | 67.40 | 80.10 |
| Phrank | 3,035 | 28.64 | 49.94 | 54.19 |
| Phen2Disease | 3,044 | 25.70 | 46.05 | 50.98 |

The earliest Exomiser trajectory group contains 157 NF1 cases from one publication among 268 cases. The corresponding LIRICAL group contains 225 cases, 70.7% from three publications. These groups show within-group trajectories, not representative temporal effects for an arbitrary patient population. Each Phrank and Phen2Disease group contains 256 ANKRD11 cases among 475 (53.9%), with 67.2% supplied by its three largest publications. Full-cohort comparisons and the shared cohort remain separate analyses.

##### Candidate presence and ranking changes

We partitioned paired cases by whether their causal gene was returned at baseline, at the endpoint, at both or at neither. A group's contribution is its sum of paired Top-1 credit differences divided by the full system cohort size, multiplied by 100. Contributions therefore sum to the overall change and are not conditional accuracy within each group.

For Exomiser, 39.36 of the 40.53 percentage-point net increase occurs among cases with the gene returned at both endpoints. Candidate entry contributes 1.17 points. In the other systems, candidate entry contributes more of the net increase, while changes among cases with the gene returned at both endpoints contribute 15.19–21.56 points.

Later databases did not improve every patient result. Fractional Top-1 credit decreased for 70 Exomiser, one LIRICAL, 122 Phrank and 73 Phen2Disease patients. These losses remain in the paired means.

Returned does not mean previously implicated in disease. Exomiser can return a gene because a variant passes filtering, even without a human disease annotation. The partition also does not imply an unchanged set of competitors or identify which annotations changed a rank. Some genes disappeared from the output without causing a Top-1 loss, because their baseline Top-1 credit was already zero.

**Table S3:** Candidate-presence transitions and contributions to the net Top-1 change. Denominators are 5,886 Exomiser, 1,090 LIRICAL, 3,062 Phrank and 3,071 Phen2Disease cases. Contributions are percentage points of whole-cohort accuracy; sums can differ by rounding. These are descriptive paired point estimates.

| Group | Exomiser | LIRICAL | Phrank | Phen2Disease |
|---|---|---|---|---|
| *Case counts* | | | | |
| Returned at both endpoints | 5,671 | 303 | 1,514 | 1,523 |
| Returned only later | 100 | 697 | 1,477 | 1,477 |
| Returned only at baseline | 3 | 0 | 1 | 1 |
| Absent at both endpoints | 112 | 90 | 70 | 70 |
| *Contribution to net Top-1 change (pp)* | | | | |
| Returned at both endpoints | 39.36 | 21.56 | 18.06 | 15.19 |
| Returned only later | 1.17 | 58.90 | 36.44 | 36.06 |
| Returned only at baseline | 0.00 | 0.00 | 0.00 | 0.00 |
| Absent at both endpoints | 0.00 | 0.00 | 0.00 | 0.00 |

#### Baseline annotation status

The main text says for how many patients the baseline release held no phenotype association for the causal gene, and what that group's gains contribute. This section defines that status, gives the counts, and lists the cases where the definition needs care.

Baseline status is read from the original files of each patient's own baseline knowledge-database release: annotated if the causal gene has at least one phenotype association there, unannotated if it has none. For Exomiser the lookup covers the human phenotype data of its native profile; its mouse, fish and interaction evidence is separate and can rank a gene with no human annotation. Beside this primary definition, the status as each system's configuration read it is reported as well. The two differ for 34 LIRICAL and 7 Exomiser patients, whose diseases are annotated only through Orphanet entries their configurations do not read, and for no Phrank or Phen2Disease patient: present in the snapshot but not read is neither a conversion loss nor curation lag.

Table S4 cross-tabulates baseline status with the final-release source classes of the audit. The two axes stay apart: a group defined at baseline and a group defined at the final release never form one paired temporal effect. Table S5 gives each group's accuracy before the case report and at the final release, and its whole-cohort contribution with 95% publication-cluster intervals. A patient whose own publication is cited by a causal-gene association, but never as its sole support, stays in the no-exclusive-support class (264, 11, 23 and 23 patients).

Exomiser's source classes here are counted on the phenotype-aspect pairs of its native human profile: 838 patients all-exclusive, 2,509 some-exclusive. Counted over every profile pair, inheritance and onset included, none is all-exclusive and the same 3,347 patients are some-exclusive. The main text's publication-support table is unaffected: its Exomiser count is the same under either denominator.

**Table S4:** Baseline annotation status by final-release source class. Counts of patients. Status is read on each patient's own baseline knowledge-database release, from its original files; the classes are the run's source audit at each system's final release. Exomiser's status and classes are read on the human phenotype data of its native profile.

| System | Baseline status | All exclusive | Some exclusive | No exclusive support | No association at final release | Total |
|---|---|---|---|---|---|---|
| Exomiser | Unannotated at baseline | 837 | 742 | 124 | 380 | 2,083 |
| | Annotated at baseline | 1 | 1,767 | 2,012 | 23 | 3,803 |
| LIRICAL | Unannotated at baseline | 494 | 206 | 9 | 15 | 724 |
| | Annotated at baseline | 0 | 306 | 60 | 0 | 366 |
| Phrank | Unannotated at baseline | 846 | 613 | 18 | 70 | 1,547 |
| | Annotated at baseline | 7 | 1,000 | 507 | 1 | 1,515 |
| Phen2Disease | Unannotated at baseline | 846 | 613 | 18 | 70 | 1,547 |
| | Annotated at baseline | 7 | 1,009 | 507 | 1 | 1,524 |

**Table S5:** Top-1 accuracy by baseline annotation status. Accuracy in percent before the case report and at each system's final knowledge-database release; contributions in percentage points of whole-cohort accuracy, with 95% publication-cluster intervals.

| System | Baseline status | Patients | Before case report (%) | Final release (%) | Contribution, pp (95% CI) |
|---|---|---|---|---|---|
| Exomiser | Unannotated at baseline | 2,083 | 5.62 | 58.91 | 18.86 (14.51–23.80) |
| | Annotated at baseline | 3,803 | 48.07 | 81.61 | 21.67 (16.22–28.20) |
| LIRICAL | Unannotated at baseline | 724 | 0.00 | 84.25 | 55.96 (42.31–70.13) |
| | Annotated at baseline | 366 | 3.01 | 75.96 | 24.50 (11.37–37.78) |
| Phrank | Unannotated at baseline | 1,547 | 0.00 | 72.13 | 36.44 (28.05–45.94) |
| | Annotated at baseline | 1,515 | 12.75 | 49.25 | 18.06 (12.63–24.11) |
| Phen2Disease | Unannotated at baseline | 1,547 | 0.00 | 71.58 | 36.06 (28.05–45.45) |
| | Annotated at baseline | 1,524 | 14.91 | 45.51 | 15.19 (9.69–21.54) |

#### Between-system comparisons on the identical patients

The shared cohort puts the same 1,090 patients through all four systems, so between-system differences can be paired patient by patient. Table S6 gives the four accuracies and the six paired differences with 95% publication-cluster intervals, at the historical baseline and at each system's final knowledge-database release, and repeats both for the 724 patients unannotated at all four baselines. Between the two knowledge states the order of the systems reverses, in both sets: Exomiser leads at the baseline and trails at the final release. At the baseline each system runs at its own, per-patient resource state: the comparison is between the systems as they were historically usable, not at one common date.

**Table S6:** Between-system comparisons on the identical patients. Top-1 accuracy and paired between-system differences with 95% publication-cluster intervals, on the shared cohort and on its patients unannotated at all four baselines. A negative difference means the first-named system trails. At the baseline each system uses its own per-patient knowledge-database release.

| | Before case report | Final release |
|---|---|---|
| *Identical patients of all four systems (1,090 patients; 108 publications)* | | |
| Accuracy (%) | | |
| Exomiser | 20.09 | 50.18 |
| LIRICAL | 1.01 | 81.47 |
| Phrank | 2.81 | 74.04 |
| Phen2Disease | 2.95 | 67.72 |
| Paired difference, pp (95% CI) | | |
| Exomiser − LIRICAL | 19.08 (8.87–31.36) | −31.28 (−45.62 to −17.11) |
| Exomiser − Phrank | 17.28 (8.30–29.36) | −23.85 (−40.84 to −8.41) |
| Exomiser − Phen2Disease | 17.14 (8.50–29.10) | −17.54 (−34.24 to −1.34) |
| LIRICAL − Phrank | −1.80 (−5.28 to 0.74) | 7.43 (−2.17 to 16.63) |
| LIRICAL − Phen2Disease | −1.94 (−5.88 to 0.72) | 13.74 (3.05–23.57) |
| Phrank − Phen2Disease | −0.15 (−1.32 to 0.88) | 6.31 (2.28–9.86) |
| *Of them, unannotated at all four baselines (724 patients; 71 publications)* | | |
| Accuracy (%) | | |
| Exomiser | 6.35 | 41.02 |
| LIRICAL | 0.00 | 84.25 |
| Phrank | 0.00 | 81.17 |
| Phen2Disease | 0.00 | 73.90 |
| Paired difference, pp (95% CI) | | |
| Exomiser − LIRICAL | 6.35 (3.97–9.09) | −43.23 (−56.36 to −28.95) |
| Exomiser − Phrank | 6.35 (3.93–9.04) | −40.15 (−53.87 to −25.81) |
| Exomiser − Phen2Disease | 6.35 (3.89–9.14) | −32.87 (−47.36 to −18.02) |
| LIRICAL − Phrank | 0.00 (0.00–0.00) | 3.08 (−5.37 to 10.88) |
| LIRICAL − Phen2Disease | 0.00 (0.00–0.00) | 10.36 (1.11–19.26) |
| Phrank − Phen2Disease | 0.00 (0.00–0.00) | 7.28 (3.37–10.99) |

#### Recent baseline releases

To assess whether large effects require very old baselines, we restricted cases to a baseline release no more than 90 days before the publication lower bound. Top-1 changes remained between 35.2 and 80.0 percentage points (Table S7). The audit also retains 30- and 180-day restrictions. These thresholds were chosen after examining the original results and are exploratory.

Across the full cohorts, median baseline-to-publication gaps were 64, 25, 30 and 32 days for Exomiser, LIRICAL, Phrank and Phen2Disease; maximum gaps were 475, 115, 115 and 127 days, respectively.

This check restricts the age of the baseline release. It does not resolve the diagnosis-to-publication interval, curation delays, phenotype vocabulary drift or source-paper overlap. Because the restricted cohort differs from the full cohort, the difference between effect sizes is not the amount explained by snapshot age.

**Table S7:** Top-1 accuracy with baseline releases within 90 days of publication. Baseline and final accuracies are percentages; changes and 95% publication-cluster confidence intervals are percentage points. Each row compares the same restricted cases at both endpoints.

| System | Cases | Baseline | Final | Change | 95% CI |
|---|---|---|---|---|---|
| Exomiser | 3,611 | 37.9 | 72.5 | 34.5 | 25.0–46.8 |
| LIRICAL | 1,053 | 0.9 | 80.9 | 80.0 | 71.8–87.1 |
| Phrank | 3,025 | 6.4 | 60.5 | 54.1 | 43.6–64.7 |
| Phen2Disease | 2,852 | 7.8 | 58.8 | 50.9 | 38.9–63.8 |

#### Recorded phenotype compatibility

Phrank and Phen2Disease record missing and mapped phenotype identifiers. Restricting to cases with no recorded missing terms at either endpoint retained Top-1 changes of 54.0 and 50.5 points, respectively. Excluding mapped identifiers as well produced similar point estimates (Table S8).

For Exomiser and LIRICAL, selected native scoring checks are reported under "Native phenotype scoring and full-output checks." They do not supply a cohort-wide historical term-mapping sensitivity comparable to the Phrank and Phen2Disease restrictions here.

These checks concern recorded identifier handling. They do not establish identical ontology semantics, rule out every representation effect or reconstruct the symptoms known at the original visit. Missing diagnostic fields for Exomiser and LIRICAL are classified as unknown, not as zero missing terms.

**Table S8:** Top-1 changes after restricting to cases with recorded phenotype compatibility at both endpoints. Changes and 95% publication-cluster intervals are percentage points. "Mapped" refers to the recorded identifier mappings; this does not audit all semantic changes in the ontology.

| System | Restriction | Cases | Change | 95% CI |
|---|---|---|---|---|
| Phrank | No missing terms | 2,998 | 54.0 | 43.2–64.7 |
| Phrank | No missing or mapped terms | 2,996 | 54.0 | 43.2–64.7 |
| Phen2Disease | No missing terms | 2,995 | 50.5 | 38.9–62.6 |
| Phen2Disease | No missing or mapped terms | 2,993 | 50.4 | 39.0–62.7 |

#### Condition concentration and patient identity

Equal weighting of genes, publications and recorded disease-label sets retained positive Top-1 changes in all four systems (Table S9). These means show that the sign of the release effect is not confined to the original patient weighting. They answer different questions and do not establish generalization to unrepresented conditions. The ten largest genes account for 26.1–37.7% of cases, depending on the system.

**Table S9:** Descriptive Top-1 changes under alternative weighting. The last four columns are percentage-point differences using equal weight per patient, gene, publication or complete recorded disease-label set. Gene and publication counts refer to the paired cohort. No new intervals are estimated for the alternative weights.

| System | Genes | Papers | Patient | Gene | Paper | Disease set |
|---|---|---|---|---|---|---|
| Exomiser | 453 | 823 | 40.5 | 52.2 | 50.2 | 52.2 |
| LIRICAL | 101 | 108 | 80.5 | 77.5 | 74.7 | 77.1 |
| Phrank | 273 | 351 | 54.5 | 66.7 | 64.4 | 66.1 |
| Phen2Disease | 273 | 352 | 51.2 | 70.3 | 66.8 | 70.0 |

The cross-publication screen flags matching gene, disease and phenotype profiles. Similarity is not proof that two records describe the same person; dissimilar descriptions do not prove independence. The current flags have not been treated as confirmed duplicates.

The finite identity audit did not establish a merge for any of its 17 candidate records. Excluding the candidates present in each system's cohort retained Top-1 increases of 40.48, 80.44, 54.56 and 51.14 points for Exomiser, LIRICAL, Phrank and Phen2Disease, respectively. This sensitivity does not establish independence of all patients or families.

Repeated genomic backgrounds provide a separate source of dependence. Linking consumed VCF hashes to frozen preparation receipts identified 255 backgrounds in the Exomiser cohort and 251 in LIRICAL. A paired bootstrap over these donors retained positive intervals (Table S10). These are separate sensitivity intervals, not a joint correction for donors and publications.

**Table S10:** Top-1 increases with 95% donor-cluster bootstrap intervals. The paired point estimates are unchanged; the resampling unit is the background donor.

| System | Patients | Donors | Change (pp) | 95% CI |
|---|---|---|---|---|
| Exomiser | 5,886 | 255 | 40.53 | 38.94–42.15 |
| LIRICAL | 1,090 | 251 | 80.46 | 78.10–82.69 |

#### Observed release transitions

The saved timeline records each case's first returned causal candidate, first positive Top-1 credit and first full Top-1 credit. Each event is accompanied by the previous evaluated release. Events already present at baseline and events never observed are recorded separately; no transition date is interpolated. Reversals are retained. The complete event tables are the run's `supporting/observed-transitions.csv`, one per system, under `main-experiment/main/` of the data deposit.

Much of each phenotype-only system's increase arrives in a single knowledge-database step (Table S11): the step into the February 2026 annotations carries 21.1, 20.2 and 46.9 points for Phrank, Phen2Disease and LIRICAL, while each system's next credit-changing step is small. Exomiser's largest step is its 2602 phenotype update (10.7 points). In LIRICAL's second row the phenotype files are byte-identical on both sides: the 19 credits gained there come from variant resources, not phenotype annotations.

**Table S11:** The knowledge-database step into February 2026 and each system's next credit-changing step. Cases are the patients comparable across the step; contributions are percentage points of whole-cohort Top-1 accuracy. A row with the same phenotype release and annotation count on both sides is a step in which another component of the data state changed.

| System | Data state | Phenotype release | Annotation count | Cases | Credit gained | Credit lost | Contribution (pp) |
|---|---|---|---|---|---|---|---|
| Exomiser | 2025-09-26 to 2026-02-28 | 2508_phenotype to 2512_phenotype | 303,574 to 284,847 | 5,868 | 61 | 23 | 0.65 |
| | 2026-02-28 to 2026-05-13 | 2512_phenotype to 2602_phenotype | 284,847 to 294,573 | 5,886 | 705 | 73 | 10.74 |
| LIRICAL | 2026-01-08 to 2026-02-16 | v2026-01-08 to v2026-02-16 | 7,492 to 7,624 | 1,072 | 515 | 1 | 46.93 |
| | 2026-02-28 to 2026-05-13 | v2026-02-16 to v2026-02-16 | 7,624 to 7,624 | 1,090 | 19 | 0 | 1.74 |
| Phrank | 2026-01-08 to 2026-02-16 | v2026-01-08 to v2026-02-16 | 322,515 to 329,339 | 3,044 | 885 | 343 | 21.12 |
| | 2026-02-16 to 2026-06-09 | v2026-02-16 to v2026-06-06 | 329,339 to 331,738 | 3,062 | 234 | 309 | 5.56 |
| Phen2Disease | 2026-01-08 to 2026-02-16 | v2026-01-08 to v2026-02-16 | 322,515 to 329,339 | 3,053 | 644 | 23 | 20.17 |
| | 2026-02-16 to 2026-06-09 | v2026-02-16 to v2026-06-06 | 329,339 to 331,738 | 3,071 | 170 | 4 | 5.41 |

#### Established associations and dating evidence

This sensitivity asks whether the aggregate changes conceal heterogeneity among associations labelled as already established. It uses the corrected patient outcomes and original evidence-basis labels; it does not require or validate the omitted first-discovery classifications. The original established category combines independent curated evidence with earlier reports in the study's source cases. The estimates differ substantially (Table S12). This evidence-basis split is not a validated clinical disease mix or a record of what each database contained. NF1 belongs to the source-case-prior group despite its longstanding disease association. LIRICAL's curated group includes 81 EHMT1 cases from one publication; excluding that publication leaves 68 cases and a 7.35-point change (interval −5.88 to 36.36). This illustrates concentration within small selected strata.

**Table S12:** Established-association results split by the original basis of dating evidence. Baseline and final accuracies are percentages; changes and 95% publication-cluster intervals are percentage points. The labels describe their original evidence basis, not a completed first-discovery classification. These intervals use 10,000 draws and the follow-up seed 20261111.

| System | Cases | Papers | Baseline | Final | Change (95% CI) |
|---|---|---|---|---|---|
| *Independent curated evidence* | | | | | |
| Exomiser | 1,387 | 80 | 83.9 | 89.9 | 6.1 (3.2–12.7) |
| LIRICAL | 149 | 18 | 6.7 | 52.3 | 45.6 (-3.6–70.5) |
| Phrank | 810 | 51 | 22.3 | 38.6 | 16.4 (3.7–37.8) |
| Phen2Disease | 819 | 52 | 25.4 | 31.2 | 5.8 (-5.9–24.8) |
| *Earlier source case* | | | | | |
| Exomiser | 1,961 | 399 | 21.0 | 72.6 | 51.6 (42.8–61.1) |
| LIRICAL | 209 | 28 | 0.0 | 90.9 | 90.9 (84.1–97.2) |
| Phrank | 707 | 111 | 1.7 | 58.2 | 56.5 (44.9–68.6) |
| Phen2Disease | 707 | 111 | 2.4 | 55.4 | 53.1 (40.6–65.8) |

### Publication sources and native scoring

#### How the four-system publication-support table was calculated

Table 1 groups patients by recorded support in the final phenotype databases. Inclusion requires at least one association linked to the causal gene whose only recorded source is the paper describing the patient. Other associations for the same gene need not have exclusive support. The table uses the corrected cohorts and saved paired outcomes; it changes neither databases nor rankings.

Historical and final Top-1 accuracies divide the included patients' summed credit by their number. A contribution to the overall increase sums their paired changes in Top-1 credit, divides by the full system cohort size and multiplies by 100, yielding percentage points. This contribution can exceed the net cohort increase if other patients lose credit. Confidence intervals use 10,000 publication-cluster resamples, retaining paired outcomes and group membership together and recomputing the full-cohort denominator in every draw. These contributions locate observed score changes; they do not estimate the causal effect of the cited findings.

The source files establish which publications support the phenotype associations used by each system. Exomiser's final native phenotype database (2602) lacks citations, so exact disease–phenotype pairs were matched to its declared upstream HPO annotation release of 8 January 2026. The other three systems use the 3 September 2026 annotations.

#### How much final accuracy occurs in each publication-support group

To relate the source audit to the reported benchmark score, we accounted for final Top-1 credit in all four publication-support groups. The calculation uses the saved rankings for every Phrank and Phen2Disease case. Table S13 reports accuracy within each group, its contribution to the full cohort's accuracy, and its share of final credit. A contribution divides the group's summed fractional credit by the full cohort size; a credit share divides it by the cohort's total final credit. Both are multiplied by 100, yielding percentage points and percentages, respectively.

The groups distinguish causal genes with all, some or no phenotype associations supported exclusively by the patient's publication, plus genes absent from the annotation file. Associations are counted across all diseases linked to the causal gene. A publication can supply patients to several groups, so publication counts are not additive. Absence of exclusive support does not establish independent knowledge. The table accounts for observed scores, not the effect of deleting annotations.

**Table S13:** Final Top-1 accuracy by recorded publication support in the 3 September 2026 annotations. Group accuracies use group sizes; contributions use the full cohort; credit shares use its total final credit. Parentheses give 95% publication-cluster intervals. Contributions and shares sum to cohort accuracy and 100%, respectively, before rounding; interval endpoints need not sum.

| Exclusive support | Patients | Papers | Final Top-1 (%) | Final contribution pp (95% CI) | Credit share % (95% CI) |
|---|---|---|---|---|---|
| *Phrank* ($n = 3,062$) | | | | | |
| All rows | 853 | 96 | 78.23 | 21.79 (15.65–29.00) | 35.84 (27.07–44.86) |
| Some rows | 1,613 | 196 | 69.95 | 36.85 (28.05–46.10) | 60.60 (51.22–69.70) |
| No rows | 525 | 53 | 12.64 | 2.17 (0.74–3.72) | 3.56 (1.12–6.97) |
| Gene absent | 71 | 9 | 0.00 | 0.00 (0.00–0.00) | 0.00 (0.00–0.00) |
| *Phen2Disease* ($n = 3,071$) | | | | | |
| All rows | 853 | 96 | 78.44 | 21.79 (15.96–28.80) | 37.15 (29.14–45.70) |
| Some rows | 1,622 | 197 | 61.94 | 32.71 (25.91–40.28) | 55.78 (46.76–64.74) |
| No rows | 525 | 53 | 24.24 | 4.14 (0.66–8.99) | 7.07 (1.10–16.49) |
| Gene absent | 71 | 9 | 0.00 | 0.00 (0.00–0.00) | 0.00 (0.00–0.00) |

For the 853 patients whose audited annotations all cite only their own publication, the contribution to final accuracy also equals the contribution to the temporal increase. Each of these patients has zero baseline Top-1 credit in the frozen results. The equality therefore holds in every bootstrap draw, giving identical endpoint and temporal contribution intervals for this group in the source data. Table 1 combines the all- and some-exclusive groups and therefore has different intervals. Credit shares require their own denominator, recomputed in each draw. The two systems contain the same patients in this group, so the findings are not independent patient replications.

The accounting can be checked directly from the supplied case-level records. We verified that all four groups sum to final accuracy and the paired temporal increase, in the observed data and every bootstrap draw. The calculation was added after inspecting the temporal results; it changes no annotations and requires no new rankings. The corrected calculation and source-classification checks are recorded in `corrected-analysis/provenance-verification.json` under the submission-closure archive. Manuscript tables are generated by `paper/results-work/integrate_corrected.py`.

The joined cases, unrounded estimates and bootstrap metadata are available in `source-data/provenance-endpoint-case-audit.csv`, `source-data/provenance-endpoint-summary.csv` and `source-data/provenance-endpoint-bootstrap-metadata.json`.

Table S14 completes the main text's publication-support table with the exact complement: the patients with no exclusively own-paper-supported association identified. Only Exomiser's complement contributes measurably to the increase (9.64 points); in the other three systems the complement's contribution is indistinguishable from zero.

**Table S14:** With and without exclusive own-publication support at the final release. The with-support rows repeat the main text's publication-support table; the complement rows are the remaining patients of each cohort. Accuracy in percent; contributions in percentage points of whole-cohort accuracy, with 95% publication-cluster intervals.

| System | Group | Patients | Before case report (%) | Final release (%) | Contribution, pp (95% CI) |
|---|---|---|---|---|---|
| Exomiser | With exclusive support | 3,347 | 21.75 | 76.07 | 30.89 (23.96–38.83) |
| | No exclusively own-paper-supported association identified | 2,539 | 47.93 | 70.28 | 9.64 (6.89–12.83) |
| LIRICAL | With exclusive support | 1,006 | 0.40 | 86.88 | 79.82 (71.52–86.83) |
| | No exclusively own-paper-supported association identified | 84 | 8.33 | 16.67 | 0.64 (−0.19 to 2.24) |
| Phrank | With exclusive support | 2,466 | 4.98 | 72.81 | 54.63 (44.23–64.94) |
| | No exclusively own-paper-supported association identified | 596 | 11.83 | 11.13 | −0.14 (−0.65 to 0.35) |
| Phen2Disease | With exclusive support | 2,475 | 3.71 | 67.63 | 51.51 (40.87–63.11) |
| | No exclusively own-paper-supported association identified | 596 | 22.69 | 21.35 | −0.26 (−1.02 to 0.35) |

#### Native phenotype scoring and full-output checks

Native replay checks whether a source-linked association reaches a scoring calculation, beyond its presence in a database. For Exomiser, the human-phenotype component was replayed on all 5,886 corrected patient inputs. Comparison with three selected complete diagnostic outputs reproduced 16 term-level components and two retained human disease-model scores without numerical disagreement. The causal gene was absent from the third output, so its component replay is not treated as confirmation of use in a complete ranking.

For LIRICAL, native replay used the observed and explicitly absent findings of all 1,090 patients. All 198 phenotype components compared with three selected complete outputs agreed exactly. These checks establish calculation parity at the stated scope; they do not test whether removing an association would change the final gene rank. For Phrank and Phen2Disease, reconstructing the native gene and disease profiles from the audited annotations reproduced the recorded input hashes. This checks that the source audit examines the profiles consumed by those ranking implementations.

The native replay and comparison records are `exomiser-components/native-verification.json`, `exomiser-components/full-trace-parity.json` and `lirical/component-verification.json` under the cohort-source archive. The Phrank and Phen2Disease profile checks are recorded in `corrected-analysis/provenance-verification.json` under the submission-closure archive.

Our Exomiser executions also match outputs we did not produce: at the one knowledge-database release shared with an independent group's public Exomiser run (Zenodo record 22967674), our execution reproduces their phenotype scores to all printed decimals for every shared patient whose causal gene was returned.

#### Example: a patient-linked association in LIRICAL scoring

A native LIRICAL trace shows how a phenotype association supported by a tested patient's paper contributes to the disease score. For `PMID_38977784_F01_II_1`, the patient's observed phenotype `HP:0000486` matched the same term in the profile for `OMIM:621182`, linked to *NAV3*. The audited assertion cites only `PMID:38977784`, the publication reporting the patient. The native calculation returned a phenotype likelihood ratio of 3.2824. A value above one favours the candidate disease, demonstrating a non-neutral scoring contribution. This does not establish whether that contribution changed the top-ranked gene or quantify the effect of the publication on overall accuracy.

The full witness table holding this assertion regenerates from public inputs with the released code; its verification record, `lirical/component-verification.json`, carries the table's counts.

#### NF1 and the Exomiser release components

The 157 NF1 cases introduced under "Release dependence and cohort concentration" have no Top-1 successes at their historical baseline and all succeed at the final endpoint. Their median rank changes from 78 to 6 at the 1802-to-1805 hg38 archive transition, with phenotype release 1807 fixed; 50 cases reach Top-1. The 1711 and 1802 variant-store hashes are identical. Between 1802 and 1805, only the variant-store member changes; the recorded genome and transcript members are identical. The two other single-archive phenotype transitions each contribute one net Top-1 credit.

Another 106 NF1 cases first reach Top-1 at the release introducing a ClinVar whitelist. This temporal coincidence does not identify the responsible records or filter decisions. Excluding the entire NF1 publication leaves a 38.92-point main Exomiser contrast (95% publication-cluster interval, 31.99–46.48). Native source traces and conversion checks are preserved separately; they do not identify a single variant-record cause for this whole trajectory. The corrected candidate-count and single-archive step tables are included in the accompanying source data.

#### Source removal: the exact rule and what it changed

The main text reports the removal in three sentences; this section carries their weight. It states the exact deletion rule, because the result depends on edge cases the short description cannot settle; it accounts for what the deletion touched; and it shows, patient by patient, how the deletion contrast relates to the dated contrast.

The rule is conservative: an association is deleted only when its support is unambiguously the tested case's publication and nothing else. Each removal targets one publication, the single PubMed identifier the patient catalogue records for the case; all patients of one publication share one removal. In the annotation file, a row counts as the publication's alone only if every one of its recorded references is that publication; rows with no reference, with references that are not PubMed identifiers or DOIs, or with any other publication among their references are kept. A disease–phenotype association is removed only when every positive row supporting it qualifies; rows recording the absence of a feature (negated annotations) never count as support. When an association is removed, every gene–phenotype row derived from it goes with it, for every gene and every disease the publication alone supports, not only the tested patient's. Gene–disease links, the ontology and frequency fields stay untouched. Phrank's model is then rebuilt from the edited file by the temporal experiment's own normalizer and the unmodified Phrank load, so candidate sets and information content are recomputed per publication. A publication that alone supports no association removes nothing. The release edited is the temporal run's final HPO state (v2026-09-01, in force from 3 September 2026); every input is pinned by SHA-256 to the identities that run recorded. Each patient is scored on the release as published and on the release edited for their publication.

What the deletion touched: of the 1,690 publications, 1,137 lose at least one gene–phenotype row (21,083 rows, carrying 21,065 associations); 1,099 change Phrank's model, and 38 lose rows while the model stays as published, because each lost pair survives through another disease of the same gene — a count of "publications whose removal changed the resource" must say which of the two it uses. The 853 cohort patients whose every causal-gene association is supported exclusively by their own publication (1,057 among all accepted patients) are exactly the patients whose causal gene leaves the candidate set; they stay in the denominators with zero credit. A further 71 cohort patients (107) have a causal gene without any association at the final release and are never ranked under either condition. Credits with rank ties are kept as exact fractions throughout; 939 cohort patients have a fractional final Top-1 credit.

The main text says the dated contrast and the deletion contrast mostly point the same way;

Table S15 is that claim in full. The dated contrast compares each patient's credit at the final release with their credit at the historical baseline. The deletion contrast compares the same final-release credit with the credit after the deletion. On each contrast a patient ends with less, the same or more credit. Each patient therefore falls into one of nine cells, and the table shows the counts at Top-1, Top-3 and Top-10, with all margins. Most patients fall in the less–less cell: their final-release credit disappears at the historical baseline, and it disappears again after the deletion. The exceptions stay visible instead of being averaged out. Patients in the less–same cell, for example, gained from later releases yet lose nothing under the deletion: their gain does not depend on associations their publication alone supports. The table counts agreement and claims nothing more; neither contrast is divided by the other.

The released removal tables also recompute the dated contrast for this pairing. The estimate equals the main analysis; the interval endpoints differ in the second decimal, because the bootstrap stream is labelled differently. The main analysis is the source to quote.

Two checks tie the execution to the main experiment. With nothing removed, every cohort patient's recomputed outcome equals the clean run's final-state result. And for each publication, the file Phrank was given was rebuilt independently from the release's provenance audit and matched by checksum: the deleted rows are exactly the rows the audit attributes to that publication alone.

**Table S15:** Directions of the two paired contrasts per cohort patient. Rows: credit at the historical baseline; columns: credit after the deletion; "less", "same" and "more" are relative to the patient's credit at the untreated final release. Panels give Top-1, Top-3 and Top-10; each sums with its margins to the 3,062 cohort patients.

| | Credit after removal | | | |
|---|---|---|---|---|
| Credit at the historical baseline | Less | Same | More | Total |
| *Top-1* | | | | |
| Less | 2,011 | 328 | 2 | 2,341 |
| Same | 24 | 575 | 0 | 599 |
| More | 1 | 121 | 0 | 122 |
| Total | 2,036 | 1,024 | 2 | 3,062 |
| *Top-3* | | | | |
| Less | 1,962 | 383 | 1 | 2,346 |
| Same | 14 | 587 | 0 | 601 |
| More | 1 | 114 | 0 | 115 |
| Total | 1,977 | 1,084 | 1 | 3,062 |
| *Top-10* | | | | |
| Less | 1,884 | 443 | 0 | 2,327 |
| Same | 10 | 603 | 0 | 613 |
| More | 1 | 121 | 0 | 122 |
| Total | 1,895 | 1,167 | 0 | 3,062 |

#### Locating the supporting records

Tables and figure data are in `paper/results-work/source-data/`. Paths beginning with `source-data/` are relative to `paper/results-work/`. Corrected analysis, patient-date and conversion records are under `data/reviewer-followup/submission-closure-20261115/`; cohort-source records are in its `cohort-source-analysis/` subdirectory. The inventory in `paper/reviewer-followup/manuscript-supplement-alignment.json` records which tables, figures and source files belong to this version. Earlier exploratory artifacts remain archived and are excluded from that inventory's current scientific artifact list. Path names containing "reviewer-followup" and "submission-closure" refer to the internal review process described in the manuscript's "Use of large language models" section, not to a prior journal submission.
